# Fast and accurate taxonomic domain assignment of short metagenomic reads using BBERT

**DOI:** 10.1101/2025.09.07.674730

**Authors:** Dmitry Alekhin, Moshe Alon, Tomer Sidi, Stav Perez Mazeh, Gon Carmi, Omri M. Finkel, Amir Erez

## Abstract

Shotgun metagenomes from complex environments such as soil uncover vast biodiversity. Yet most short reads produced by shotgun sequencing cannot be taxonomically or functionally annotated, as they lack a sufficiently comprehensive reference, obscuring the true structure and function of microbial communities. We introduce BBERT, a nucleotide large language model optimized for short reads. Testing on a large cohort of soil metagenomes, we found that BBERT identifies bacterial sequence syntax without relying on reference databases, enabling accurate assignment of taxonomic domain, coding potential, and reading frame directly from reads as short as 100 bp. BBERT is small and fast enough to analyze metagenomes using a modest GPU and can be used to convert short metagenomic reads directly to bacterial amino acid sequences for downstream applications. BBERT also improves de-novo metagenomic assembly, reducing mismatches and gaps while accelerating runtime. Using metagenomes from wild legume nodules, we demonstrate that BBERT filtering improves bin quality while significantly accelerating de-novo assembly. By providing fast, reference-free classification of short reads, BBERT unlocks large metagenomic archives for more accurate ecological and evolutionary analyses.

**Graphical Abstract:** 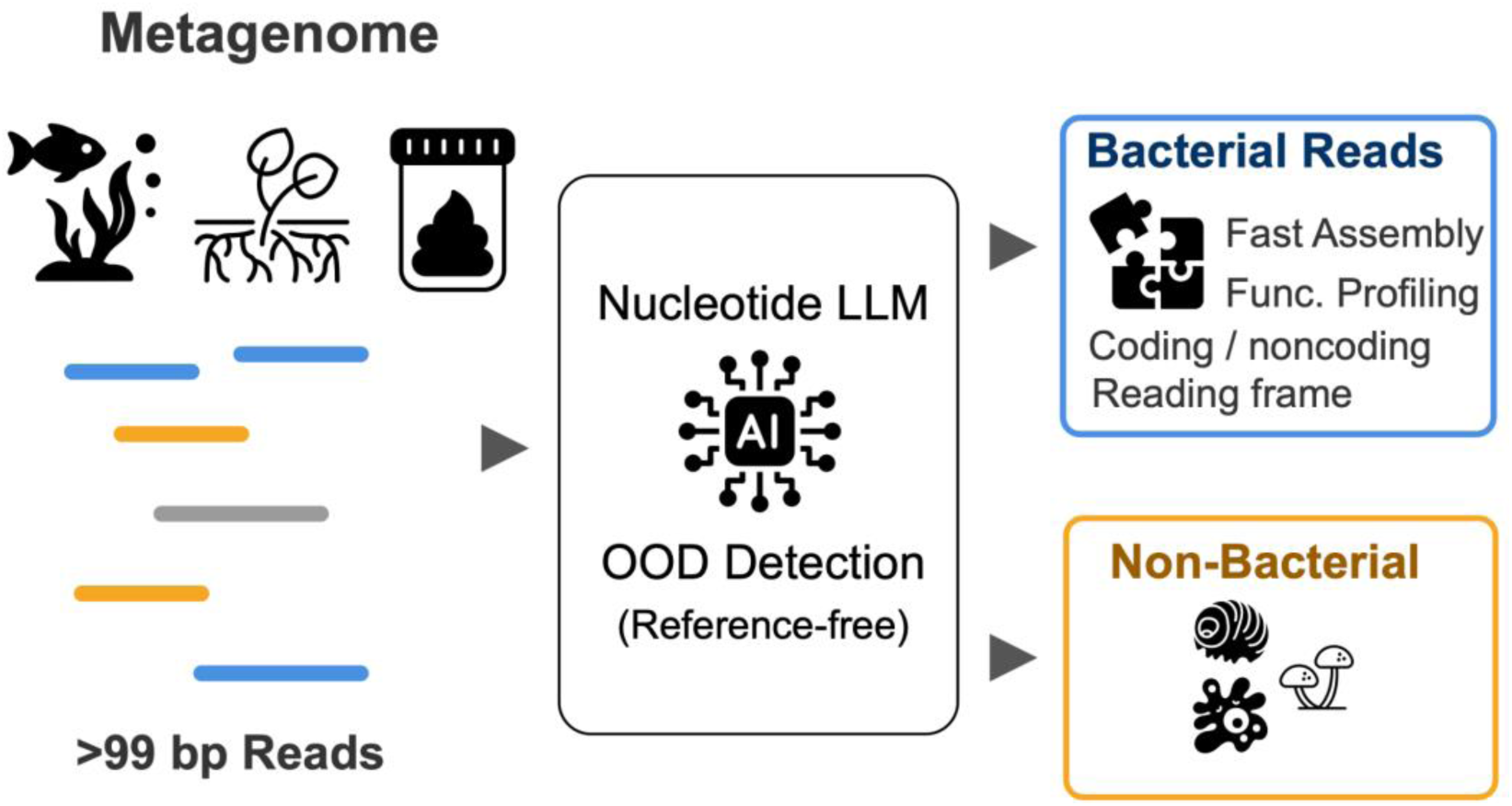

## Introduction

Microbial ecosystems encompass immense complexity. Understanding this complexity is essential for elucidating the processes that sustain the biosphere, a task that relies on the analysis of massive sequence datasets. Today, such datasets are abundant: thousands of publicly available metagenomes, spanning every biome on Earth, can be freely downloaded. Among these, soil metagenomes are extraordinarily important for us to understand. Soil is the substrate on which all terrestrial life, including humanity, thrives. Yet, despite centuries of study, we still have only a partial understanding of what makes a healthy soil (1). Soil habitats contain diverse representations of all domains of life: Bacteria, Archaea, Eukaryota, and their viruses.

Since many species found in soil are absent from any reference database (2–5), current annotation tools are unable to taxonomically classify many, or in some cases most, of the DNA reads even at the domain level. This phenomenon is commonly referred to as “microbial dark matter” (6, 7). The partiality of the databases forces existing bioinformatic tools to employ heuristics to classify metagenomic reads (8, 9) which causes many of the reads to be either unannotated or mis-annotated. Notably, a significant and variable portion of this “dark matter” in soil metagenomes appears to be non-bacterial, much of it eukaryotic in origin (10–12). This causes a *conflation problem*: a comparable measurement of normalized functional profiles from soil metagenomes requires correctly identifying and separating eukaryotic and bacterial DNA reads within a given metagenome. However, no method currently exists that can reliably solve the conflation problem for short reads. Therefore, it is not possible to correctly normalize bacterial gene abundances and obtain a comparable set of values between different systems. The conflation problem is exacerbated by the vast difference in genome structure and size between prokaryotes and eukaryotes. Without a reliable domain-level classifier, neither the function-specific read counts (numerator) nor the appropriate total read count defining the background (denominator) can be determined for functional abundance estimation (10–12).

Beyond taxonomic classification, the functional annotation of short metagenomic reads faces additional serious difficulties. In the absence of comprehensive reference databases, it is hard to determine which reads code for proteins and what is their reading frame. This *annotation problem* means unknown proteins that exist in the data remain unknown. Additionally, even when the protein is known, the need to check all six reading frames of each read slows the analysis while inflating false discoveries. Finally, the conflation problem also causes an *assembly problem*. De-novo assembly of short metagenomic reads into contigs is highly resource-demanding and prone to errors. It makes sense that by separating the reads to taxonomic domains, the assembly problem could be ameliorated.

To address the aforementioned conflation, annotation and assembly problems, we developed BBERT: a BERT-based (13, 14) large language model (LLM) that answers three simple questions: is a given metagenomic read bacterial in origin? Is it protein coding? If so, what is the reading frame? To overcome the sparsity of non-bacterial reference genomes, we trained BBERT using an out-of-distribution (OOD) detection approach, flagging reads that differ from the learned bacterial patterns as non-bacterial. By focusing on the primary divergence in the tree of life and training only on bacterial reads, we were able to achieve high accuracy and speed even with a modestly-sized model. Thus, BBERT overcomes both throughput and accuracy limitations of previous LLM-based methods (15–17). Using BBERT, we were able to perform large-scale analyses essential for the exploration of soil metagenomes. We partitioned the bacterial coding reads from within thousands of soil metagenomes. We further demonstrated that BBERT prefiltering improved annotation speeds and the quality of de-novo assemblies of metagenomic data, reducing errors and improving bin quality.

## Materials and methods

BBERT is a BERT-based model developed for out of distribution (OOD) detection of nucleotide sequences originating from bacterial genomes: we trained on bacterial sequences to detect whether a sequence is bacterial or not. Here, we outline a detailed description on BBERT’s training and evaluation (Methods Section 1). Then, the design of two synthetic scenarios used to implement BBERT within the context of downstream tasks, namely frame classification and metagenome assembly (Methods Section 2). Finally, we describe the methodological approach for applying BBERT to complex environmental datasets, with a focus on soil metagenomes (Methods Section 3).

### 1. BBERT training and evaluation

To create a nucleotide-based LLM for OOD detection of bacterial genomes, we adopted three phases of development: (i) pretraining the BERT architecture only on bacterial genomic reads, (ii) fine-tuning for the task of bacterial/non-bacterial classification, and (iii) evaluation on the bacterial read classification task with comparison to BERTax (15), Tiara (16) and REMME (17).

BBERT pretraining was performed on a bacterial dataset of sampled synthetic short nucleotide sequences, each 100 bp in length, designed to simulate metagenome reads and their errors (see *Datasets* and *Preprocessing* sections). For the training process, we used the Masked Language Model (MLM) objective, i.e. masking 30% of tokens in the input and optimizing the model’s ability to infer those tokens from context, which enables the model to learn representation of missing nucleotides based on those reads. Notably, we use the average cross entropy loss calculated on the masked nucleotides only. The pretraining optimization process was conducted in sessions of 5 to 10 epochs until an optimal result was achieved, saving all intermediate checkpoints (model weights). For each checkpoint, we evaluated the classification accuracy on the calibration dataset (see the *Datasets* section) and selected the optimal checkpoint accordingly. Typically, we observed simultaneous decreases in training loss and increases in classification accuracy. Training was terminated when progress stagnated, with no obvious improvement in model performance.

Next, to use BBERT for bacterial read classification, we attached a lightweight classification head: a linear layer trained on top of the BBERT encoder’s token embeddings, to directly predict whether a read is bacterial or not. The head was trained in a supervised manner on the same calibration dataset described in the *Datasets* section, with equal numbers of bacterial and eukaryotic species and no taxonomic overlap at the genus level to the pre-training dataset.

Rather than keeping the pretrained BBERT weights frozen, we fine-tuned the encoder jointly with the classification head, allowing its representations to adapt to the classification objective. The head and encoder were optimized with separate learning rates (1e-5 and 1e-6, respectively, AdamW optimizer) to avoid destabilizing the pretrained weights. The resulting fine-tuned encoder was subsequently used as a fixed feature extractor for reading-frame prediction and coding/non-coding classification, with only the corresponding task-specific prediction heads being trained.

To evaluate the model performance, we constructed another 10 disjoint datasets, with no taxonomic overlap at the genus level with the calibration or the pretraining datasets, namely the evaluation datasets. We calculated each read score and classified it as bacterial according to a probability cutoff of 50% (bact_prob > 0.5), reporting the overall classification metrics on those datasets. The results on this benchmark were compared to BERTax (15), Tiara (16), and REMME (17). We further compared the encoder fine-tuning strategies described above against the fully frozen baseline. Fine-tuning the encoder substantially improved classification performance: read-pair accuracy on the held-out evaluation set increased from 0.869 (frozen encoder) to 0.899 with the best partial-unfreezing configuration, with corresponding improvements in F1 score (0.869 to 0.900), and AUROC (0.937 to 0.959). Among the configurations tested, freezing only the bottom two transformer layers (closest to the input embeddings) while fine-tuning all higher layers achieved the best overall performance. We therefore adopted this configuration as the default BBERT classifier throughout the remainder of this work.

#### Accelerating and training REMME

Although REMME is a state-of-the-art DNA language model, it required inference optimization and adaptation for bacterial/non-bacterial classification to enable a fair comparison with BBERT. REMME is pretrained using three prediction objectives: masked-token reconstruction, reading-frame classification, and coding-fraction regression. For the downstream tasks considered in this work, however, only the task-specific predictions are required. We therefore streamlined the inference pipeline by removing computation of the masked-token reconstruction head and replacing row-wise prediction aggregation with vectorized operations. Together, these optimizations reduced inference time by 10-50-fold compared with the original analysis pipeline, without changing the downstream predictions.

Bacterial/non-bacterial classification is not one of REMME’s original pretraining objectives. In the original work (17), the pretrained REMME encoder is adapted to a downstream task by fine-tuning it together with a task-specific prediction head (REBEAN for enzymatic function prediction). Following the same adaptation strategy, we attached a three-layer feed-forward classification head to the pooled representation at the first sequence position (the same representation used by REMME’s native frame-classification head) and jointly fine-tuned the encoder and the new classifier, with no frozen layers, on labeled bacterial/non-bacterial read data.

To benchmark REMME against BBERT across all three tasks (reading-frame classification, coding classification, and bacterial-origin classification), we built matching inference and evaluation pipelines that run BBERT’s own task-specific heads (separately trained for each of the three tasks) on the same test datasets, using identical evaluation metrics, confusion-matrix analyses, and inference-throughput measurements for both models.

We did not retrain REMME for reading-frame prediction because the published model predicts only four coarse frame classes (three reading-frame offsets and a non-coding class), whereas BBERT predicts the full six-class signed reading frame (and separately coding/noncoding). To enable the fairest possible comparison, we derived the optimal one-to-one mapping between REMME’s predicted frame classes and the ground-truth labels before computing frame-classification accuracy, thereby giving REMME the most favorable alignment for this evaluation.

#### Datasets

For BBERT development and evaluation we curated two types of datasets. One for the pretraining, that comprises 1416 bacterial species (for a good representation of the bacterial tree-of-life, we randomly sampled two genomes from each bacterial family in the RefSeq database), and an additional dataset for calibration and evaluation that comprises 899 bacterial species from 396 different genera and 1200 eukaryotic species from 823 different genera.

Notably, the two datasets are separated at the genus-level and share no genus among them. Additionally, as we compare the model performance with BERTax (15), we verified that the latter does not share any genus with the published BERTax training and finetuning sets.

REMME’s pretraining strategy makes its input genomes less transparent and therefore it is possible that some genera used for REMME’s pretraining are present in our test datasets, giving it potentially some advantage; nevertheless, BBERT performs better than REMME (see *Results*). The pretraining genome accession list is provided in Table S5.

To construct a curated reference set of soil-relevant eukaryote genomes, we first established a target taxon list by reviewing 18S rRNA and ITS amplicon metabarcoding studies of soil communities, including the Earth Microbiome Project (18) and the pan-European LUCAS soil survey (19), as well as global surveys of soil protist (20, 21) and fungal diversity (22, 23). From this literature synthesis, eight major eukaryotic lineages consistently detected as abundant members of soil communities were identified: *Cercozoa*, *Ciliophora*, *Oomycota* (Heterokonts), *Glomeromycota* (arbuscular mycorrhizal fungi), *Mucoromycota*, *Ascomycota*, free-living *Nematoda*, and *Amoebozoa*, supplemented by soil-associated *Chlorophyta* and terrestrial plant taxa with sequenced genomes. Within each lineage, genera were selected based on their reported prevalence in soil metabarcoding datasets and the availability of sequenced genomes in public repositories.

To account for noise in species group selection, all accessions from the calibration and evaluation dataset were divided separately by genus into 20 equal groups for bacteria and 20 equal groups for eukaryotes, with random abundances assigned following a log-normal distribution within each group. These groups were then merged pairwise to form 20 combined subsets, each with a unique genus composition from both domains. From these, 10 subsets (datasets 1–10) were assigned for fine-tuning, while the remaining 10 (datasets 11–20) were reserved for evaluation. Species abundance files for score calibration and evaluation sets (group 1–10 for score calibration, groups 11–20 for evaluation) can be found in Table S6.

#### Dataset preprocessing

BBERT’s focus is to classify metagenomic reads: short nucleotide sequences that may contain errors. These errors typically include insertions, substitutions, and deletions in DNA sequences. Thus, to train and test BBERT on data resembling typical metagenomic sequencing, we generated synthetic short reads with typical sequencer errors. The genomes were downloaded from the NCBI database (in .fna format), and using InSilicoSeq (24), we simulated reads from each .fna file using the NextSeq error model.

#### BBERT model and pretraining

BBERT is based on the Hugging Face implementation of the BERT architecture (BertForMaskedLM). During training, the data collation process plays a critical role in preparing raw DNA reads into consistent, model-ready batches. Specifically, the collation function performs token masking, label assignment, and sequence padding, ensuring that all inputs in a batch are of uniform length and properly annotated for training loss computation. To accommodate for nucleotide sequence input, we implemented a simple character-based tokenizer, with five core tokens: A, C, T, G, and N. The latter, N, represents ambiguous or error-prone bases. To support sequence structure and model training, we added special tokens: <s> (start-read),</s> (end-read), <pad> (padding), and <msk> (mask for the masked language modeling, MLM, objective). Reserved tokens like <cls> (read summary), <sep> (read separator for multi-read inputs), and <unk> (unknown nucleotide) were also included for extended flexibility. During training, only reads with a length of at least 100 nucleotides were used, truncating longer reads to a fixed length of 100 tokens. Then, 30% of the tokens in each read were randomly selected and replaced with <msk>, while the remaining tokens were assigned a label of -100 to exclude them from the loss computation. The model was trained to predict the original identity of the masked tokens based solely on their surrounding context, using a cross-entropy loss computed over the masked positions. This self-supervised learning strategy allows BERT to capture the underlying structure and dependencies within DNA sequences, enabling effective downstream classification. Details of the dataset generation strategy are provided in the <u>Datasets</u> section.

#### Model inference

This section outlines the procedure for applying BBERT to new metagenomic read datasets. The scoring pipeline supports common input formats, including *FASTA*, *FASTQ*, and compressed *gzip* files. During preprocessing, each read was truncated to 100 characters, all letters were converted to uppercase, and any character not belonging to the canonical nucleotide set [A, C, T, G] was replaced with ‘N’. Reads were tokenized using the same vocabulary employed during training. For each batch, a per-read score was computed using the cross-entropy loss function, consistent with the training objective.

To improve classification robustness in paired-end sequencing data, the inference pipeline processes both R1 and R2 reads independently, computing classification scores for each. A separate script can then be used to compute the mean of the two as the final score for the read pair. If only one of the R1 or R2 reads is 100 bp or longer, we use its score directly without averaging, to maintain reliability. The data are processed in batches (default batch size = 1024 reads) for efficient computation within the model. For scalability, particularly when handling large input files, the data are divided into larger chunks, each containing 50 times the batch size by default. This chunking strategy ensures efficient CPU memory usage and enables parallel processing.

#### Hardware and software

All model training procedures were carried out on an NVIDIA A100 GPU with 80 GB of VRAM, using a batch size selected to maximize efficient resource utilization and model performance.

For inference and benchmarking, models were converted from 32-bit (FP32) to 16-bit floating-point precision (FP16) to achieve faster scoring, with a minimal trade-off in accuracy. On our evaluation datasets (11–20), the accuracy loss due to the transition from FP32 to FP16 was less than 0.1%. A further performance increase stems from batch inference; the user can choose the batch size most suitable for their hardware.

All benchmarks were conducted across different hardware configurations to evaluate performance under diverse computational environments: a high-performance computing cluster node equipped with an NVIDIA A100 GPU (80 GB VRAM), optionally utilizing a multiprocessing DataLoader; cluster nodes with NVIDIA L40 (48GB VRAM) and NVIDIA L4 (24GB VRAM); a local workstation featuring an NVIDIA RTX 3090 GPU (24 GB VRAM) and 64 GB of RAM; and a cloud-based Google Colab instance running an NVIDIA Tesla T4 GPU (16 GB VRAM). This range of setups allowed us to assess the model’s efficiency from powerful cluster nodes to accessible cloud resources.

#### Throughput benchmark

To assess the computational performance of BBERT, we conducted a comparative benchmark against existing approaches, BERTax (15), Tiara (16) and our custom-fine-tuned version of REMME (17). The evaluation was performed under optimized conditions for each model, using the maximum batch size that fully utilized the available GPU memory.

### 2. Synthetic data: using BBERT for important downstream tasks

#### Reading frame classification

#### Single case visual analysis – Low dimensional representation of the BBERT embedding space

Synthetic 150 bp reads were generated using a modified version of InSilicoSeq (24) for two organisms: *Bradyrhizobium diazoefficiens* USDA 110 (RefSeq ID: GCF_001642675.1) and *Saccharomyces cerevisiae* S288C (RefSeq ID: GCF_000146045.2). The simulator appended to each read’s description the start and end coordinates, as well as the percentage of coding sequence (CDS) nucleotides, computed using the corresponding .gtf file from NCBI.

BBERT was then run on the synthetic dataset to produce embeddings. Dimensionality reduction was performed using PCA and t-SNE (with t-SNE parameters: perplexity = 30, n_iter = 1000, random_state = 42), and the results were visualized using scatter plots. For PCA, an additional distribution plot was included.

#### Robust linear classification of sequence properties

To classify multiple properties of DNA reads, we trained three task-specific prediction heads on BBERT embeddings. As described above, the encoder was first partially fine-tuned for bacterial/non-bacterial classification together with the corresponding classification head. The resulting encoder was then kept frozen and used as the feature extractor for the remaining tasks. Separate linear classification heads were trained independently for (i) six-class reading-frame prediction and (ii) coding versus non-coding classification. Thus, bacterial/non-bacterial classification employed partial encoder fine-tuning, whereas the reading-frame and coding classifiers evaluated the representation learned by the fine-tuned encoder without further modifying it.

For bacterial/non-bacterial classification, we constructed the training dataset by combining the first 10 mixed calibration sets (Datasets 1–10), while evaluation was performed on the remaining 10 mixed subsets (Datasets 11–20), ensuring no overlap in genus composition between training and test sets.

(i) The training dataset for frame classification was constructed using the same accession numbers from the NCBI database as those used for training the BBERT model; however, only coding regions were included. From each coding region, a fragment of around 300 base pairs was randomly extracted, with minor variation in length introduced by sampling from a normal distribution with a mean of 300 bp and standard deviation of 5 bp. Reads of 100 bp length were then taken from the beginning and end of this fragment. To simulate sequencing errors, artificial mutations, insertions, and deletions were introduced using the error matrices from InSilicoSeq(24). This procedure resulted in a dataset comprising 10M reads, which was used to train the frame classification model. The test dataset was generated in the same way, but based exclusively on accession numbers reserved for BBERT evaluation dataset (see <u>Datasets</u>).

The model achieved an accuracy of approximately 95% on the evaluation datasets. The confusion matrix (see *Results*) showed that of the few misclassifications, most occurred between forward and reverse orientations, while errors between adjacent frames were rare. These results suggest that the BERT-based embeddings capture both the direction and frame position of DNA reads with high fidelity.

(ii) For the task of distinguishing coding from non-coding sequences, we used the same synthetic dataset, which was generated with the modified InSilicoSeq pipeline and includes coding percentage information in the read annotations. As in the bacterial vs. non-bacterial classification task, input reads of length 100 bp were first encoded with the frozen BBERT model to obtain contextual embeddings. These embeddings were then passed to a lightweight classifier trained to assign class 1 if more than 50% of the sequence was coding and class 0 otherwise.

#### Reduced assembly errors and run-time acceleration

To compare de-novo assembly of metagenomes before and after BBERT filtering, a synthetic dataset was generated using a modified version of InSilicoSeq (24), which was adapted to add to the read annotation the details: start, end, chromosome, organism source and % coding using NCBI’s *gtf*. Four datasets were created by composing 128 non-overlapping prokaryotic and 128 eukaryotic genomes each. Genomes were downloaded from NCBI RefSeq, and had assembly-levels of either “chromosome” or “complete genome” and were unique at the genus level. Paired-end FASTQ files (R1 and R2) were simulated using the modified InSilicoSeq. Each organism contributed 35, 000 paired-end reads. The resulting FASTQ files were mixed into combined R1 and R2 files per dataset, maintaining consistent order, to simulate a realistic metagenomic dataset. The so-called “mixed” data contain both eukaryotic and bacterial reads in equal abundance.

The BBERT model was then applied to the mixed data, generating classification scores. Based on these scores and the recommended predefined cut-off, reads were classified as bacterial or non-bacterial. We used metaSPAdes (25) to perform genome assembly on the full mixed dataset as well as on each of the classified subsets. Assembly run times were recorded, and the resulting contigs were saved in FASTA format.

To compare the assembly results, a BLAST nucleotide database was created from the 512 source genomes using the command *makeblastdb -dbtype nucl*. Each contig was aligned against this database using *blastn* with the following parameters: -outfmt 6 -max_target_seqs 10 -evalue 1e-5. Alignment errors were categorized into three types based on their position and frequency: single errors, isolated mismatches or gaps flanked by correctly aligned nucleotides; adjacent pairs, two consecutive mismatches or gaps between correctly aligned nucleotides; and longer runs, more than two consecutive mismatches or gaps.

For each organism, assembly coverage was calculated for both the mixed and classified assemblies using the formula:

(number of correctly aligned nucleotides in contigs - number of errors [mismatches or gaps]) / genome length × 100.

To further assess assembly quality, we used MetaQUAST (26) to compare the unfiltered mixed assemblies and the BBERT-filtered bacterial assemblies against the dataset-matched bacterial reference genomes. Only contigs of at least 100 bp were included. MetaQUAST was run with ‘-- min-contig 100’, ‘--unaligned-part-size 500’, and ‘--report-all-metrics’; HTML, Icarus, and plot generation were disabled. The main analysis used a minimum alignment identity of 95% together with ‘--use-all-alignments’ and ‘--ambiguity-usage all’, and a second, more permissive sensitivity analysis used a minimum alignment identity of 80% with the same alignment and ambiguity settings.

Because the aim was to quantify bacterial assembly signal and residual non-bacterial or poorly aligned sequence, MetaQUAST alignment-detail files were retained and parsed at the contig level. For each assembly, contig bases were decomposed into bacterial-aligned bases, fully unaligned contig bases, terminal unaligned bases, and internal unaligned blocks. We then compared the unfiltered and BBERT-filtered assemblies by calculating the total assembly length, bacterial-aligned sequence, unaligned sequence components, numbers of fully or partially unaligned contigs, contigs containing internal unaligned blocks, and contigs aligning to more than one bacterial reference genome.

#### Gene Ontology (GO) enrichment analysis of the synthetic data

To characterize functional biases as a function of classification outcome, we performed two GO enrichment analyses: bacterial coding reads classified as bacterial (True Positives; TP) versus bacterial reads not classified as bacterial (False Negatives; FN), and eukaryotic coding reads correctly classified as nonbacterial (True Negatives; TN) versus eukaryotic reads misclassified as bacterial (False Positives; FP). We used the four synthetic mixed bacterial/eukaryotic datasets described in Methods Section 2.b, each containing 128 bacterial genera, and restricted this analysis to bacterial coding reads. Analyses were performed separately for each comparison and for the three GO sub-ontologies: Biological Process, Cellular Component, and Molecular Function.

Because GO annotation is structured as a directed acyclic graph (DAG) following a hierarchy of resolution, annotations were propagated to higher-level terms to enable level-wise comparisons at progressively coarser resolutions. Specifically, for each annotated read, we incremented the count of its directly assigned GO term and all higher-level terms along all DAG paths, thereby generating cumulatively aggregated counts. We then constructed five “level cuts” per ontology by collecting propagated counts from the root level down through five successive depths. This procedure yielded 30 ontology-depth panels total: 15 bacterial TP/FN panels and 15 eukaryotic TN/FP panels (3 ontologies x 5 levels), each represented by a contingency table of per-term correct-classification and error-classification counts.

For eukaryotic reads, GO annotations were assigned through read coordinates, overlapping CDS features, resolved NCBI GeneIDs, and assembly-specific NCBI GAF files; organisms without a valid assembly-specific GAF were excluded from both TN and FP counting rather than being treated as having no annotated function.

For each GO term within each ontology-depth panel, we tested for differential representation using Fisher’s exact test applied to a 2×2 table comparing the correct-classification group with the error-classification group: TP versus FN for bacterial reads, and TN versus FP for eukaryotic reads. The null hypothesis assumed no association between classification outcome and membership in the GO term beyond the overall correct/error group totals within that specific ontology-depth panel. Two-sided p-values were computed, and multiple testing correction was applied within each analysis using the Benjamini-Hochberg procedure to control the false discovery rate (FDR). Effect sizes were summarized as odds ratios and normalized log_2_ fold changes, with the correct-classification group as the numerator: TP/FN for bacteria and TN/FP for eukaryotes.

### 3. Soil metagenomic analyses

#### Metagenome dataset

To find relevant shotgun-sequenced metagenomes we used the Integrated Microbial Genomes (IMG) system website. We searched for the term “soil” to get all records with this term in their descriptions. We then looked through each record to check for relevant SRA run accessions, and to verify that it was not an amplicon sequencing run. The SRA accessions were then used to acquire their respective metadata from the NCBI website. The SRA accessions for the 1969 soil metagenomes, as well as relevant metadata, can be found in SI Table S1.

#### Geographic information systems (GIS)

Metagenomes that had longitude and latitude coordinates available in their metadata were imported into ArcGIS Pro.

#### Kraken2

We used Kraken2 (27) with the following reference databases: (i) Kraken2’s expanded database with protozoa, fungi and plants (PlusPFP version September 2024) in stringent settings (confidence 0.1) as well as less stringent conditions (confidence 0). (ii) A curated Kraken2 database: we curated 213, 194 bacterial genomes downloaded from the NCBI RefSeq database. Out of all available bacterial genomes, we removed known human pathogens which represent the bulk of the database but are not relevant for our purposes. In addition, we acquired 9, 143 high-quality prokaryote genomes from the GEMs database for a total of 222, 337 genomes (SI Table S2). Reads were mapped to this database in stringent settings (confidence 0.1) as well as less stringent conditions (confidence 0), similarly to the PlusPFP database.

#### Read mapping and KEGG functional annotation

Reads were quality trimmed and mapped to the KEGG prokaryote database using DIAMOND (28) blastx with parameters --sensitive --max-target-seqs 1 --evalue 1e-5 --outfmt 6. We then assigned a KO number to read hits if available in the KEGG database for that reference hit (some reference proteins exist in the KEGG database without a corresponding KO number). We then summed up the coverage of all hits per KO in each metagenome and divided by the protein length to get an approximate number of copies of the gene in that sample.

#### Taxonomy-based MDS map of metagenomes before and after BBERT

To compare the taxonomic profile of the metagenomic samples before and after filtering by BBERT, we first used SingleM to get the taxonomic profile of each individual sample. We then combined them into one feature table that included the taxonomic profile of each sample before and after BBERT and used *metaMDS* and *vegdist* functions from the *vegan* (29) R package to calculate MDS scores, and *ggplot* to create the ordination plot.

#### Classification of organelle and eukaryotic single-copy genes by BBERT

Because mitochondria and plastids are endosymbiotic descendants of alpha-proteobacteria and cyanobacteria respectively, and because the most conserved housekeeping genes share deep homology across all domains, there is a concern that reads from these sources could be mis-classified as bacterial and artificially inflate the bacterial pool and, more specifically, the single-copy-marker pool on which our analysis relies. To address this, we simulated a mock metagenome with InSilicoSeq from the protein-coding sequences of five groups of ∼50 source genomes each, designed so that every simulated read traces back to a named source gene: (i) diverse bacteria spanning 14 phyla (positive control); (ii) complete mitochondrial genomes across metazoa, fungi, plants, algae and protists; (iii) complete chloroplast/plastid genomes across land plants, algae and diatoms; (iv) eukaryotic nuclear single-copy genes (BUSCO (30) eukaryota_odb10), the operational stand-in for “conserved housekeeping genes with deep homology across all domains“; and (v) random non-single-copy nuclear CDS from the same eukaryotic genomes (negative control). We then scored all 4.90 M reads with BBERT and report the fraction called bacterial at bact_prob ≥ 0.5 (and, stringently, ≥ 0.9).

## Results

### Reference-based tools cannot classify most metagenomic reads at the domain level

Existing methods for analyzing soil metagenomes often include de-novo assembly and binning of contiguous regions (contigs) (31). By placing metagenomic reads into longer genomic context, the classification of reads mapped to contigs becomes an easier task. Examples for such assembly-based tools include EukRep (32) and EukFinder (33), all of which perform well on data that had been assembled. Indeed, the inclusion of metagenomically-assembled genomes has vastly expanded the microbial diversity in reference databases (34). But assembly-based approaches consider only what the algorithms are able to assemble, and this represents the minority of the data. We tested this assertion by assembling 478 soil metagenomes and found that about 67%±16% of the read data could not be mapped to any of the contigs (<u>Fig. S1</u>). This demonstrates that assembly-based tools ignore most of the biodiversity in soils.

Assembly-free methods attempt to directly classify individual metagenomic reads, allowing us to gain information from all available sequence data. Such approaches include direct alignment to reference protein databases (e.g., using DIAMOND (28) on the Kyoto Encyclopedia of Genes and Genomes KEGG (35), or NCBI’s non-redundant protein database *nr* (36)), k-mer matching (Kraken2 (27)), and the use of marker genes that provide only taxonomic classification (MetaPhlAn (37)). While these methods have proven instrumental for metagenomic interpretation, their potential is capped by their dependence on external reference databases.

Approaches that attempt to transcend the limitations of reference databases include the use of hidden Markov models (HMMs), that attempt to find probabilistic features in a collection of similar reads. However, even state-of-the-art HMM-based tools such as SingleM (38), that uses HMMs for single-copy phylogenetic marker genes, do not classify the vast majority of the reads, as they do not contain such markers. Thus, while SingleM may reliably estimate bacterial composition and the fraction of bacterial reads within a sample, it cannot classify all the individual reads, limiting its use to bulk averages.

To gauge the ability of existing tools to reliably partition metagenomes at the domain level, we curated a global soil dataset of publicly available metagenomes, featuring 1969 bulk soil samples (Fig.1A and SI Table S1). We then used two of the aforementioned computational tools to measure the domain-level nucleotide fractions within each of the 1969 samples. First, we used Kraken2 (27), which uses k-mer frequencies, matched to its reference dataset. Using Kraken2’s default standard database plus protozoa fungi and plants (‘plusPFP’) resulted in an average prokaryote fraction of 15.2% ± 10.5% with an unidentified reads fraction of 82.6% ± 12.4% of the data, and only 0.1% ± 0.2% of the reads classified as eukaryotic (<u>SI Fig. S2A</u>).

**Fig. 1:**
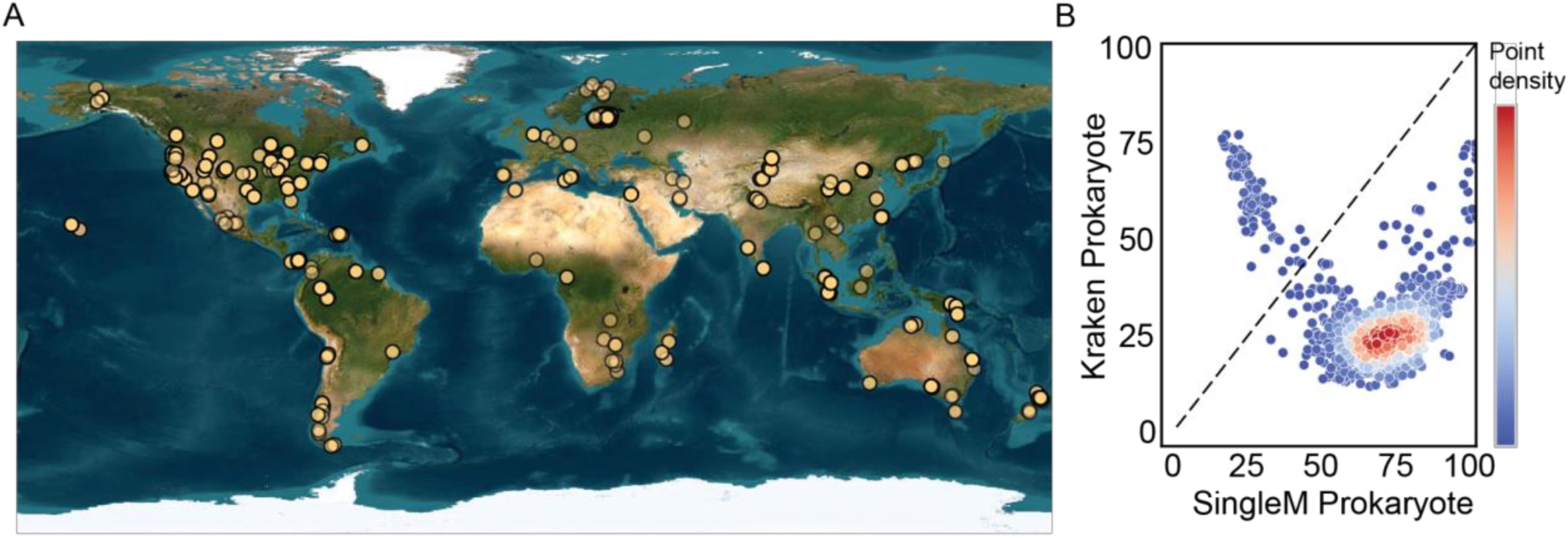
Existing state-of-the-art tools show conflicting prokaryote fractions. **(A)** The metagenomic dataset used in this study and its global distribution, compiled from 1969 soil samples. **(B)** The imputed bacterial percent of each sample, calculated using SingleM (38) on the *x*-axis and Kraken2 (27) on the *y*-axis using our custom 222, 337 prokaryote database, shown color-coded according to the local density.

Since these results are not plausible, we reasoned that the default Kraken2 database is unsuitable for analyzing soil metagenomes. In order to customize Kraken2 to the soil metagenomes we used here, we curated a custom reference database for it, consisting of 222, 337 (SI Table S2) prokaryotic isolate and metagenomically-assembled genomes (MAGs), removing commonly sequenced clinically-relevant taxa which heavily skew the composition of the genome database. Since the current database coverage of eukaryote genomes is grossly insufficient (currently in NCBI, <9, 000 reference genomes, overwhelmingly from animals, plants and fungi, out of millions of species), we did not include eukaryotes in our custom database, and simply used Kraken2 to estimate the number of prokaryotic reads in each sample. To validate the Kraken2 estimates, we used SingleM (38, 39), which uses pre-trained profile hidden Markov models (pHMM) of universal single-copy genes. Although SingleM does not classify all individual reads, but only the small fraction of reads that contain single-copy genes, it can be used to estimate the prokaryotic fraction within a metagenome. Had Kraken2 and SingleM yielded similar results, we would have concluded that Kraken2 is a practical solution for assigning taxonomy to soil metagenomic reads. We therefore compared the estimated prokaryote fraction generated by both tools (Fig. 1B and <u>SI Fig. S2B</u>). As is apparent, the two tools strongly disagree, thereby exposing a gap in our ability to accurately partition metagenomes at the domain level.

### BBERT performs consistently well on data it was not trained on

LLMs have recently emerged as powerful tools for classifying metagenomic reads by capturing complex patterns in sequence data without relying on reference databases (40, 41). Models like Evo2 (42), BERTax (15), DeepMicroClass (43) and Tiara (16), have used deep learning to improve taxonomic classification of long reads. However, these tools either perform inaccurately and inefficiently on short reads (SI Table S3), or not at all, as in the case of DeepMicroClass which has a 500 bp lower limit. This is a serious limitation, since of the approximately 100 petabytes of sequencing bases currently in the SRA archive, about 20 petabytes are from metagenome samples (44), and almost all existing metagenomic data, and most of the metagenomic data generated today, are composed of short reads, at most 300 bp long, with much of the earlier data comprising 100-150 bp reads.

LLMs inherently capture complex dependencies within DNA sequences via their attention mechanisms, enabling them to generalize beyond the training data and to perform reference-free classification of all reads. Such functionality is particularly valuable for identifying sequences absent from the training data, as LLMs can discern patterns characteristic of novel or previously unrepresented organisms (17, 45). However, current deep learning-based tools for DNA sequence analysis are prohibitively slow when applied to large numbers of short metagenomic reads (100-250 bp), restricting their use to small datasets or to long-read data (15, 42). This computational inefficiency substantially limits their utility for the large-scale metagenomic analyses needed to comprehensively explore soil biodiversity.

To overcome the limitations of existing LLM-based tools, we developed BBERT, an LLM built on Google’s BERT architecture, that can quickly and accurately classify all short metagenomic reads without relying on reference databases. BERT is based on a self-attention mechanism (13, 46), which considers the importance of each token (here, nucleotide) according to its context, and thus is well suited for DNA-read classification. DNA sequencing yields bidirectional fragments whose absolute genomic coordinates are unknown; BERT’s bidirectional encoding therefore captures both upstream and downstream nucleotide dependencies. By exploiting these capabilities, BBERT enables efficient, large-scale classification of short metagenomic reads without reliance on extensive reference databases. Since the current database coverage for bacterial genomes is much more complete than for the eukaryotic-archaeal branches of the tree of life (47), we adopted an OOD detection approach by training exclusively on bacterial sequences. The OOD framework is designed to distill the universal properties of the bacterial sequences. BBERT was trained and optimized for short reads, requiring only 100 bp to reach a conclusion quickly, opening up enormous archives.

BBERT identifies reads that do not conform to the learned bacterial sequence patterns, flagging them as non-bacterial. To this end, we pre-trained BBERT to infer masked nucleotides within reads, thus making it sensitive to the nucleotide context rather than pre-defined sequence patterns (see Methods). For the pre-training, we randomly selected two genomes out of every bacterial family from the ∼200K bacterial genomes we curated (where possible. Some families only include one sequenced genome, which we included), for a total of 1416 genomes, and used InSilicoSeq (24) to create a training set of short metagenomic reads with simulated sequencer errors (see methods and SI Table S5). We then fine-tuned the BBERT weights to classify bacterial vs. nonbacterial sequences (see *Methods*).

### BBERT embeddings correctly classify domain, coding vs. non-coding reads and identify open reading frames

LLMs operate within a high-dimensional latent embedding space; upon classification, BBERT projects each read into this representation. We hypothesized that BBERT’s high-dimensional embedding space encodes, beyond the bacteria/non-bacteria split, additional interpretable features, which may manifest as distinct clusters. Such clusters were observed in other LLM-based DNA classifiers (48, 49). To test this, we generated a mock metagenomic dataset of synthetic sequencing reads derived from two reference genomes commonly found in soil (see Methods), *Saccharomyces cerevisiae* (eukaryote) and *Bradyrhizobium diazoefficiens* (bacterium). We then inspected the 39, 168-dimensional space (size of hidden layer 384 × 102 bases including start and stop tokens) using Principal Component Analysis (PCA, Fig. 2A). As is apparent, PC1 separated the bacterial from eukaryotic reads, and moreover, the bacterial coding vs. noncoding reads. To visualize non-linear relationships in BBERT’s embedding space, we applied t-distributed stochastic neighbor embedding (t-SNE (50)), labeling reads as bacterial or non-bacterial and as protein-coding or non-coding according to their predominant nucleotide composition. t-SNE clearly separated the bacterial and non-bacterial reads, and furthermore, within each group, separated the protein coding and non-coding reads (Fig. 2B). Among the coding reads, there appeared six distinct clusters for each species. We hypothesized that these six clusters represent the six reading frames in protein-coding data. Therefore, we randomly selected several reads from each frame and labeled them. The different frames mapped almost perfectly to the different protein-coding clusters (Fig. 2C). The clear frame-wise separation in the t-SNE visualization of the embeddings led us to train a reading-frame classifier on the BBERT embeddings, which classified reads accurately (Fig. 2D). To visualize BBERT’s capability to predict open reading frames (ORFs), we ran it on a 100 bp sliding window along 20, 000 bp of the *Salmonella enterica LT2* genome, and compared BBERT’s coding probability along this window to the ORF demarcation found in NCBI’s GenBank file. We observed a high level of agreement between the two when setting the coding probability threshold to 0.75: specificity 0.96, precision 0.99, recall 0.90 (Fig. 2E). Loci where BBERT predicted a high coding probability while designated non-coding in NCBI may represent short, hitherto unidentified ORFs. These clustering patterns generalize beyond the two species examined in Fig. 2: embeddings from additional taxa exhibit analogous taxon-taxon separation and within-genome substructure (<u>Fig. S3</u>). BBERT outputs per-read embeddings (see User Manual on GitHub), enabling users to reproduce these analyses or interrogate their own datasets with standard dimensional-reduction and clustering workflows that exploit this biologically interpretable latent space. Furthermore, BBERT can therefore be used to convert short metagenomic reads to amino acid sequences for coding bacterial reads, thereby accelerating and improving many downstream applications.

**Fig. 2.**
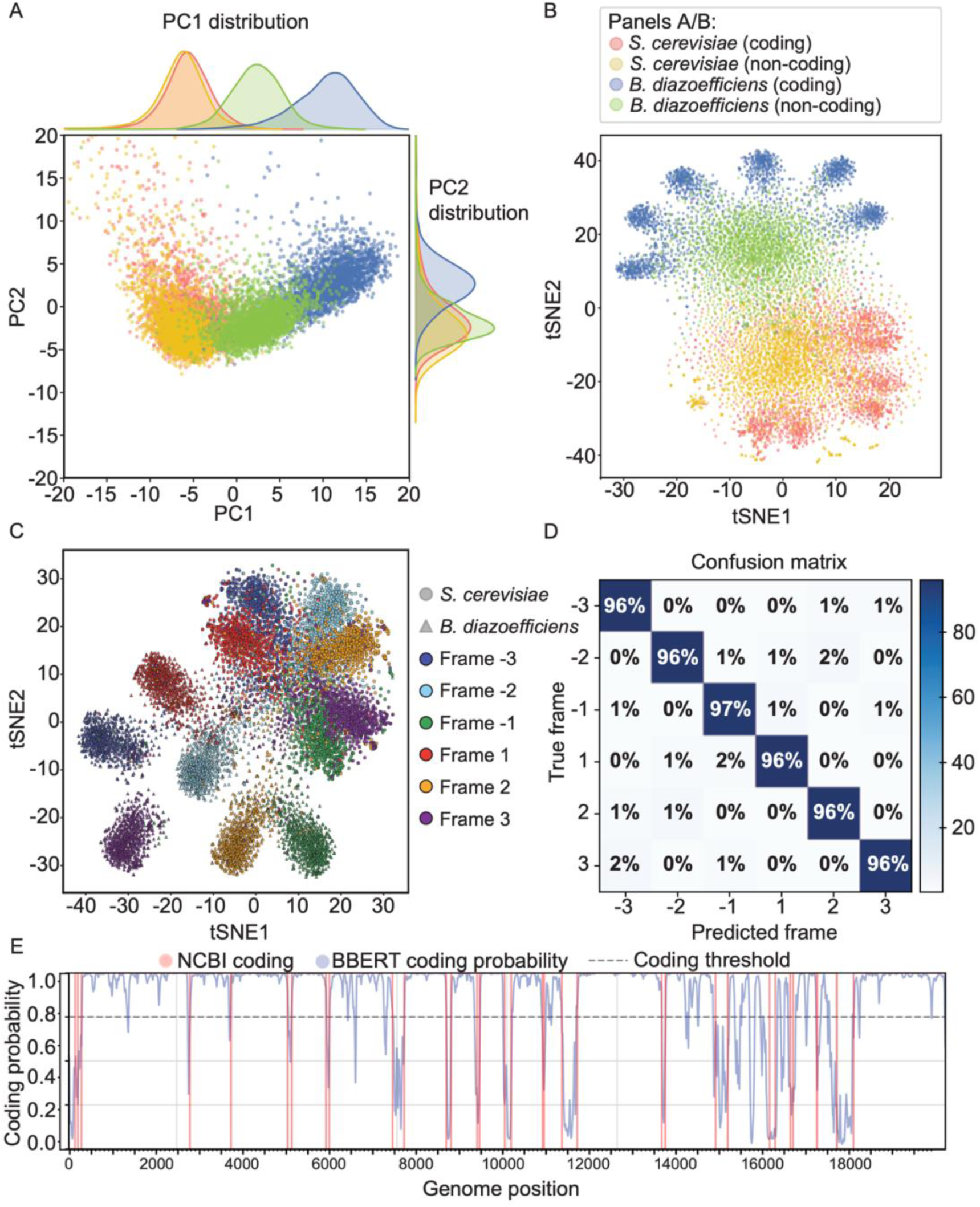
BBERT differentiates between bacteria and non-bacteria, coding and non-coding reads and identifies the reading frame. Two-dimensional visualization of the 39, 168-dimensional (102 tokens x 384 dimensions each) embedding by BBERT of short reads generated from *Bradyrhizobium diazoefficiens* (blue/green) and *Saccharomyces cerevisiae* (orange/yellow) using **(A)** principal component analysis (PCA) and **(B)** t-SNE (50). The colors represent the coding and non-coding reads. **(C)** t-SNE representation of the embedding by BBERT of the same reads as in **(B)**, labeling several reads from different reading frames (colors) and from the two species (circles and triangles), and plotting their location in the embedding space. **(D)** Confusion matrix showing BBERT’s reading frame classification accuracy on a test dataset of 100 bp reads taken from within annotated ORFs from RefSeq genomes (see SI Table S8). **(E)** NCBI ORF prediction on the *Salmonella enterica LT2* genome compared with BBERT coding probability (100 bp window).

### BBERT outperforms state-of-the-art deep-learning classifiers

The distinct clustering we observed in the embedding space (Fig. 2A-C) suggests that it would be useful to train a classifier on the BBERT embeddings. We therefore trained three such classifiers, used for determining: (i) bacterial vs. non-bacterial; (ii) coding vs. non-coding; (iii) open reading frame. We compared BBERT’s performance to two state-of-the-art deep-learning classification tools: BERTax (15) and Tiara (16). Another recent classification tool, DeepMicroClass (43), only accepts reads that are 500 bp long, and so could not be fairly evaluated. A recent tool, REMME (17), does not have a bacterial-nonbacterial classifier but we followed their manuscript to fine-tune REMME to this task. In the process of fine-tuning REMME, we realized that its run time could be significantly improved by patching their code, which we also did (see *Methods*). BBERT outperformed BERTax, Tiara and our customized REMME in standard classification metrics, including accuracy, precision, recall (on par with REMME) and AUROC. (Fig. 3A-D, SI Table S3). BBERT’s mean accuracy for domain classification on the 10 test sets is 89.7% ± 1.7%, compared with 86.8% ± 1.7% for REMME, 79.2% ± 4.7% for BERTax and 72.6% ± 3.4% for Tiara. For identifying coding reads and their reading frame, BBERT outperformed REMME, the only comparable tool that had this functionality (Fig. 3E, coding accuracy: 88.1% ± 0.9% for BBERT vs. 74.4% ± 2.1% for REMME; Fig. 3F, reading frame accuracy: 96.3% ± 0.3% for BBERT vs. 45.4% ± 0.4% for REMME).

**Fig. 3.**
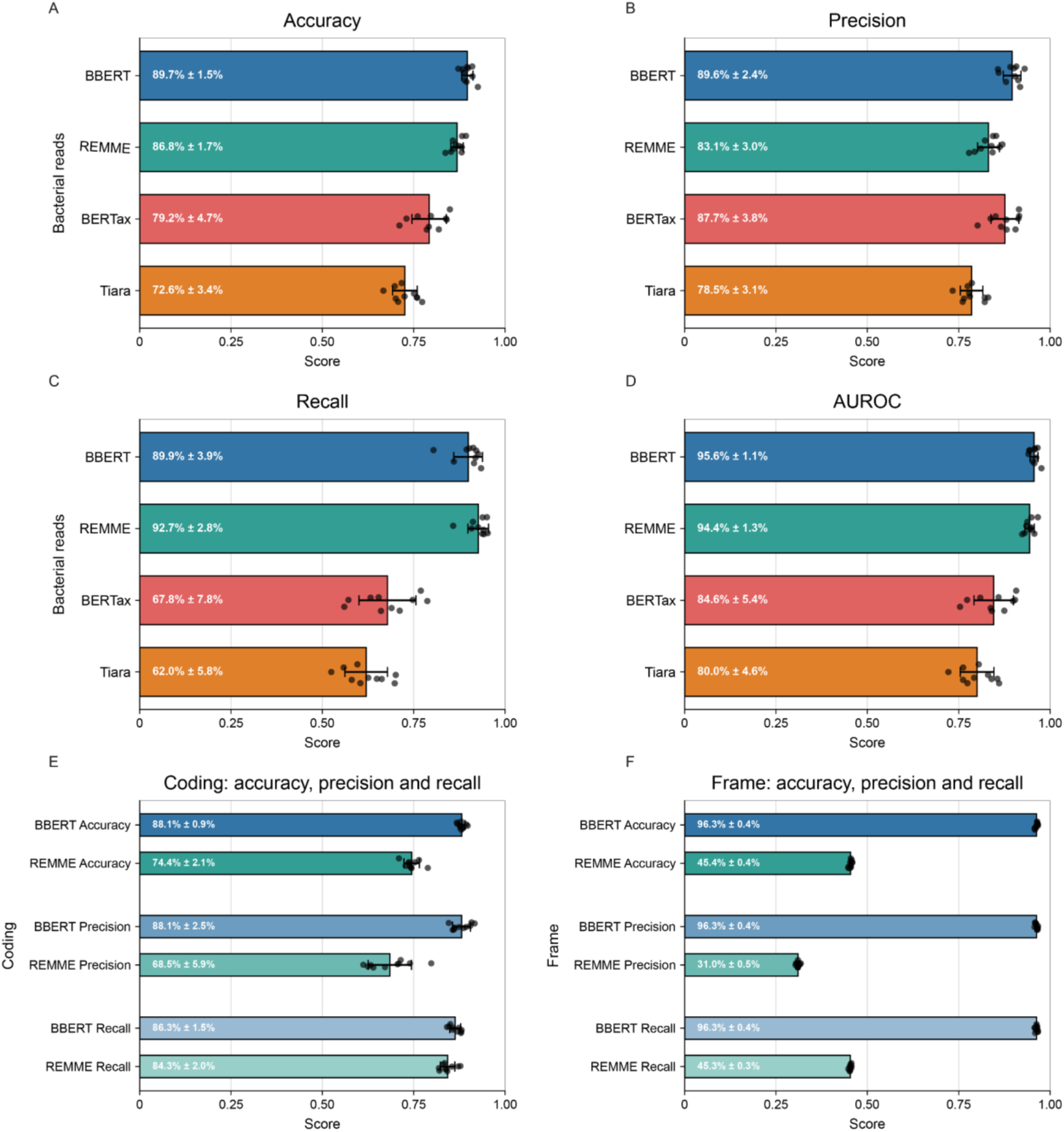
Comparison of classification accuracy. **(A-D)**, We ran the bacterial/non-bacterial classification using BBERT on the embeddings, BERTax, Tiara and our fine-tuned version of REMME. All runs were on the same test dataset (SI Table S6). **(A)** Accuracy, **(B)** Precision, **(C)** Recall, **(D)** AUROC. **(E-F)** Comparison of BBERT to REMME for **(E)** Coding/noncoding read identification; **(F)** Reading frame identification of coding reads.

Speed-wise, BBERT processed 45, 912 reads per second (rps) when running on a dedicated NVIDIA H200 GPU node, compared to 413 rps for BERTax and 347 rps for Tiara, though after patching REMME it was slightly faster at 50, 745 rps. Running BBERT on a Windows 10 personal computer with an NVIDIA RTX 3090 GPU card clocked 15, 373 rps, meaning it is usable without a high-performance computing cluster, a feat unfeasible with most alternatives (BERTax: 238 rps, Tiara: 546 rps); again, our customized REMME was slightly faster with 17, 188 rps. Indeed, one may run BBERT freely on a Google Colab T4 machine and still be able to process a typically sized metagenome in the time window this service is given for free.

Furthermore, BBERT ran on a MacBook Pro M5 Max at 3, 370 rps, meaning one can use a laptop to analyze a metagenome in a few hours. Taken together, these results demonstrate that BBERT is a fast and accurate tool to partition large amounts of short metagenomic reads at the domain level, including the large fraction of unannotated or unassembled reads.

### BBERT sorts soil metagenomic reads without taxonomic bias and agrees with SingleM estimates

We applied BBERT to our curated set of 1969 publicly available soil metagenomes. To further validate BBERT on these real-world data where the ground truth is unknown, we used the number of prokaryotic bases estimated by SingleM as a reference. We compared this SingleM estimate with (i) the total number of bases in the unfiltered metagenome; (ii) the number of bases deduced as bacterial by BBERT. Whereas before filtering, the metagenomes contained 31% more bases than deemed prokaryotic by SingleM (slope 1.31 ± 0.01), after BBERT filtering, this number was reduced to 9% (slope 1.09 ± 0.01, Fig. 4A). SingleM, however, estimates the prokaryotic fraction only using single copy genes without assigning each read individually, in contrast to BBERT which classifies all reads. In contrast with SingleM, a per-read comparison is possible against Kraken2, so we directly compared BBERT to Kraken2. As expected, BBERT and Kraken2 agree on reads Kraken2 identified as bacterial; however, BBERT additionally identifies a substantial fraction of reads that Kraken2 leaves unclassified (Fig. 4B). Conversely, there are only very few reads that Kraken2 identified and BBERT didn’t. Thus, BBERT is a sensitive bacterial identifier, allowing read-based classification like Kraken2 while unhindered by incomplete reference.

**Fig. 4.**
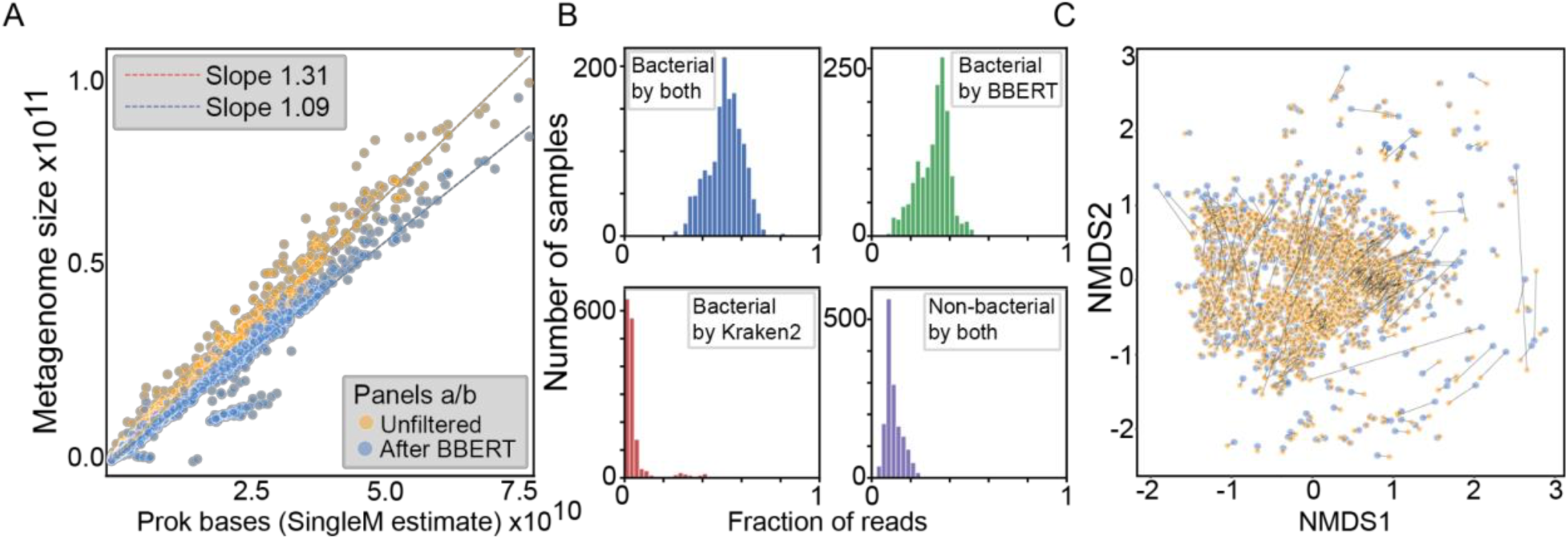
BBERT does not bias bacterial taxonomic composition and identifies the bacterial DNA fraction in agreement with SingleM. **(A)** Analysis of all 1969 soil samples reveals a striking agreement between the estimated number of prokaryotic nucleotides (by SingleM) to the actual nucleotides classified by BBERT as bacterial (blue, slope 1.09) whereas the unfiltered data contains 31% more nucleotides (orange). **(B)** Histograms displaying the fraction of reads in each metagenomic sample that were classified as (i) bacterial by both BBERT and Kraken2 with our custom database; (ii) bacterial only by BBERT; (iii) bacterial only by Kraken2; (iv) non-bacterial by both. **(C)** MDS plot of the family-level taxonomic composition, as inferred by SingleM, before and after BBERT filtering. Each sample is displayed as two dots, unfiltered and after BBERT, with a line connecting them.

To verify that BBERT filtering is not taxonomically biased, we used SingleM to profile the community composition of all metagenomes, at the family level, before and after filtering. Remarkably, with the exception of a handful of metagenomes, this composition remains nearly unchanged after BBERT filtering, demonstrating that BBERT does not bias by taxonomy (Fig. 4C). However, a small subset of samples exhibited substantially larger displacements. The two highest displacement values were recorded for hyperarid Atacama Desert soil and coastal eastern Chinese soil. A single study contributed three of the top ten samples, all replicates of one acidic urban-farm compost system. Generally, the samples most perturbed by BBERT filtering are soils with atypical or low-biomass microbial communities in desert or arid soils and acidic compost.

### Reads that fail to map to KEGG in soil metagenomes are enriched for non-bacterial sequences

Soil metagenomes are notoriously difficult to functionally annotate (3, 6–8, 23), with only a fraction of total data reliably annotated. We hypothesized that the unannotated metagenomic fraction, the so-called “dark matter”, is enriched with non-bacterial reads compared with the mapped reads. To test this, we mapped the BBERT-assigned bacterial and non-bacterial fractions from all 1969 metagenomes to the KEGG prokaryotic database, and found that at an E-value cutoff of 10^−5^, 69.7% ± 8.5% of the bacterial reads were functionally assigned, compared to 35.3% ± 10.1% in the non-bacterial fraction. This trend persists for a range of E-value cutoffs (Fig. 5A). When the percentage of reads mapped is plotted against -log₁₀(E), the data exhibit a linear relationship. The slope of this regression may serve as an indicator of read-to-database similarity, and the steeper slope observed for BBERT-sorted bacterial reads indicates a closer correspondence with KEGG. Next, we asked, out of the KEGG-mapped and - unmapped, what is the fraction of bacterial reads. For the KEGG-unmapped reads (Fig. 5B) most reads are bacterial. Nevertheless, among the KEGG-mapped reads, the BBERT bacterial fraction is much higher (Fig. 5C). Therefore, the unmapped reads are enriched with non-bacterial reads compared with the mapped reads. This remains true across a range of KEGG mapping thresholds and BBERT bacterial/nonbacterial thresholds. In summary, BBERT-based bacterial read selection increases the fraction of soil metagenomic reads that can be assigned to KEGG functions, with KEGG-unmapped “dark matter” relatively enriched for non-bacterial sequences.

**Fig. 5.**
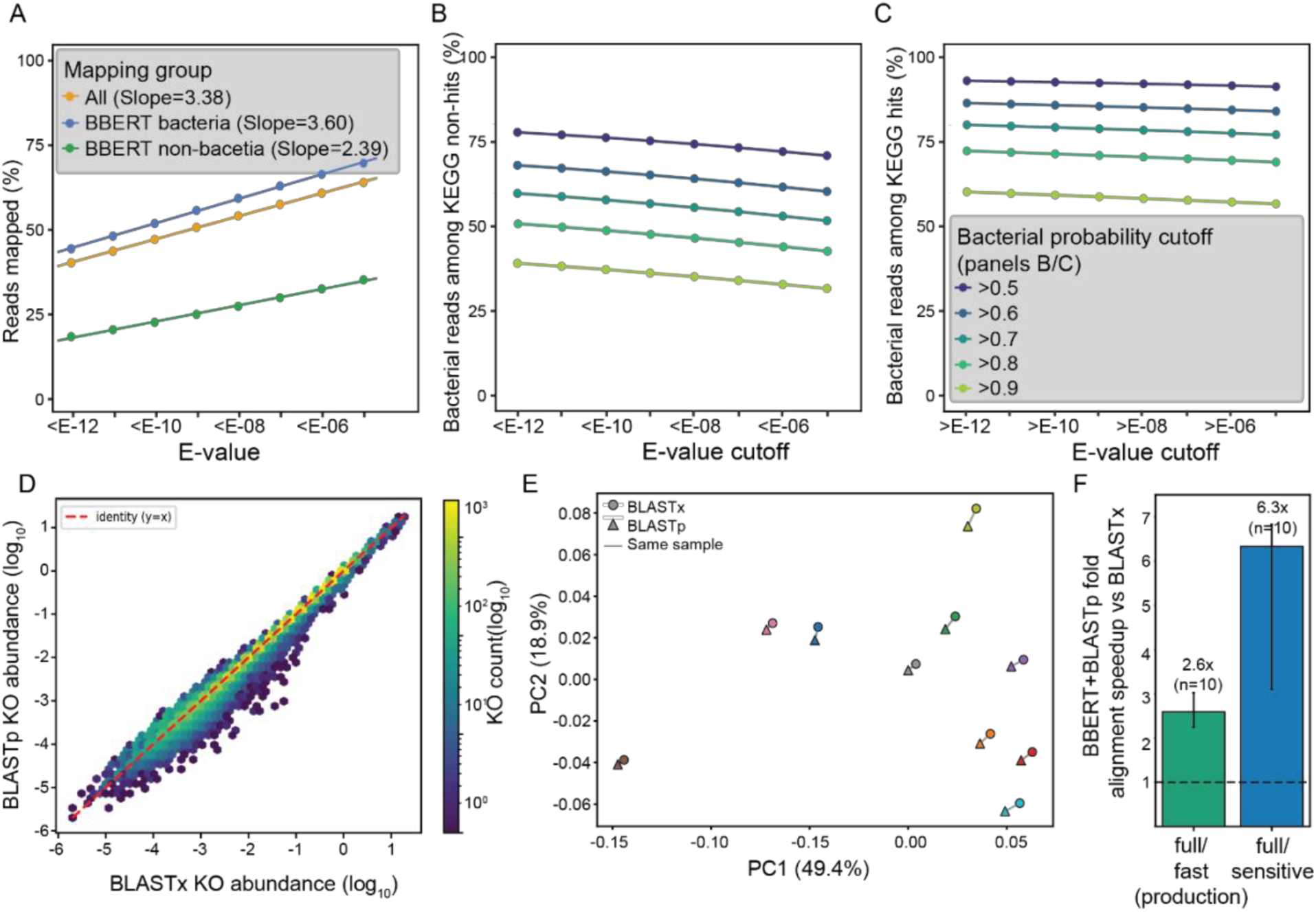
Functional implications of BBERT-sorting the bacterial reads in soil metagenomes. **(A)** Percentage of the reads mapped to KEGG prokaryote proteins from all soil metagenomes in our dataset. The values (dots) are shown for all reads (orange), and the reads sorted by BBERT as bacterial (blue) and non-bacterial (green). The dots are fitted by a straight line with the slopes reported on the figure. **(B)** Percentage of BBERT bacterial-labeled reads from KEGG non-hits defined by a BLAST E-value greater than the x-axis value. **(C)** Percentage of bacterial-labeled reads from KEGG hits defined by a BLAST E-value smaller than the x-axis value. The colors in B, C signify the BBERT bacterial probability cutoff to define a bacterial hit. **(D)** Comparison between DIAMOND BLASTp on the BBERT translated reads on the KEGG prokaryote database (*y-*axis), vs. DIAMOND BLASTx on the unfiltered reads (*x*-axis) resulted in essentially the same mapping. **(E)** PCA on the mapping results from **(D)** shows the close pairing between the filtered and translated reads with BLASTp and the unfiltered reads with BLASTx. **(F)** Run time for BLASTp vs. BLASTx shows 2.6-fold speedup in “fast” mode and 6.3-fold speedup in “sensitive” mode.

### BBERT directly translates short DNA reads into amino acids, for fast and accurate annotation

Given that BBERT identifies bacterial coding reads and their reading frame, it also directly translates short metagenomic reads into amino acid sequences. We found that running DIAMOND BLASTp on the BBERT-translated reads on the KEGG prokaryote database, resulted in essentially the same mapping as running DIAMOND BLASTx on the unfiltered reads (Fig. 5D-E). However, as expected, there is a significant speedup when using the BBERT-translated sequences, especially when running the mapping in ‘sensitive’ mode (Fig. 5F). This means that BBERT can be applied to streamline metagenomic data analysis pipelines, enabling direct queries to the reference protein database.

### BBERT accelerates de-novo metagenomic assembly with fewer errors

Could BBERT be used to improve de-novo assembly of metagenomic reads? Pre-assembly read partitioning has previously been used to reduce metagenomic assembly complexity and improve recovery of low-abundance bacterial genomes, as exemplified by Latent Strain Analysis (51) and Jorg (52). Since eukaryotic DNA is notoriously difficult to assemble, and de-novo assembly of bacterial metagenomes is a difficult task even without eukaryotic contamination, we reasoned that using BBERT to select only the bacterial reads could reduce assembly complexity and error burden while retaining most of the bacterial sequence. To test this, we created four non-overlapping synthetic metagenomes with a mix of short reads from eukaryotic and bacterial genomes (see Methods), and assembled them before and after BBERT filtering using metaSPAdes (25).

We found that BBERT filtering retained 95.6% of the unweighted mean bacterial-genome coverage obtained with the unfiltered reads (65.22% versus 68.23%; a decrease of 3.02 percentage points, or 4.4% relative; Fig. 6A, SI Table S7), consistent with the loss of some bacterial reads classified as nonbacterial. However, this reduction in coverage was not uniform across genomes. For 431 of the 512 bacterial genomes, less than 5% of the reference genome was covered only by the unfiltered assembly (Fig. 6B, SI Table S7).

**Fig. 6.**
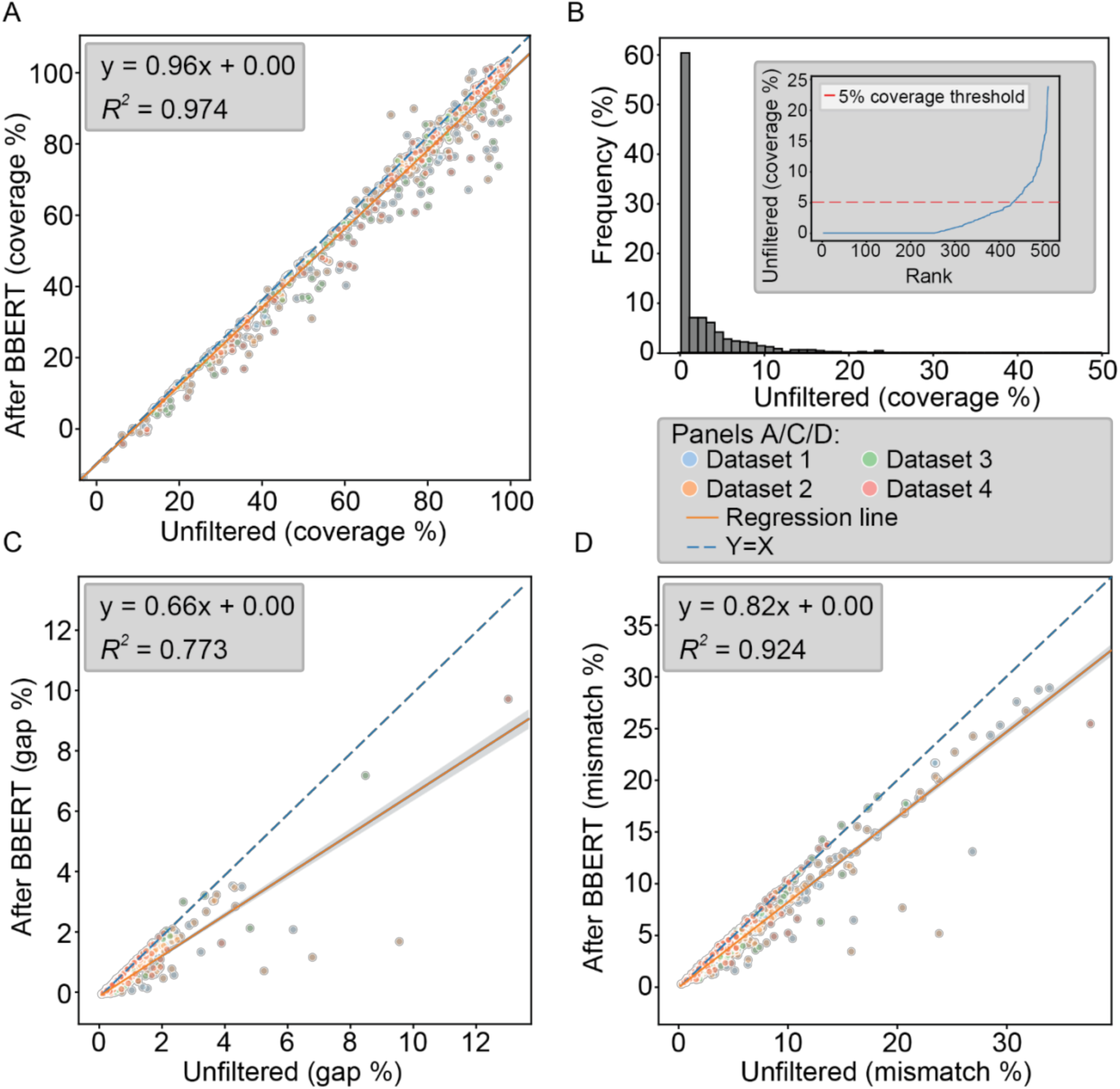
BBERT-based filtering improves de novo assembly on synthetic datasets. **(A)** Genome coverage of assembled contigs before and after BBERT classification. Each dot represents one of 128 bacterial genomes in each of four datasets (512 genomes total). The orange line is the filtered-versus-unfiltered regression through the origin (slope = 0.96), and the dashed blue line indicates *y* = *x*. **(B)** Histogram of the percentage of each reference genome covered exclusively by the unfiltered assembly. Inset: the same data ranked by unfiltered-only coverage; 431 of the 512 genomes had less than 5% of the reference genome covered exclusively by the unfiltered assembly. **(C)** Gap-opening rates normalized by covered reference sequence in unfiltered versus filtered assemblies. The filtered-versus-unfiltered regression through the origin has a slope of 0.66. **(D)** Mismatch rates normalized by covered reference sequence in unfiltered versus filtered assemblies. The corresponding regression slope is 0.82.

Moreover, the 4.4% relative reduction in mean coverage was accompanied by two practical benefits: (i) run time was reduced 2.7-fold, from 66±8 hours to 25±2 hours per metagenome, which is beneficial since de-novo assembly is an extremely resource-intensive process; (ii) regressions through the origin of rates normalized by covered sequence yielded filtered-versus-unfiltered slopes of 0.66 for gap openings (Fig. 6C) and 0.82 for mismatches (Fig. 6D). These slopes summarize the relationship across genomes. For comparison, when calculated as unweighted averages across the 512 genomes, with each measure normalized by reference-genome length (SI Table S7), gap openings decreased from 0.51 to 0.36 per 100 reference bp (a decrease of 0.15 per 100 reference bp and a 29.3% relative reduction), while mismatches decreased from 3.31 to 2.64 per 100 reference bp (a decrease of 0.67 per 100 reference bp and a 20.2% relative reduction). The combined mismatch-plus-gap-opening measure decreased from 3.82 to 3.00 per 100 reference bp (a decrease of 0.82 per 100 reference bp and a 21.4% relative reduction). These measures were lower after filtering for 499, 493, and 496 of the 512 genomes, respectively. Representative genomic windows illustrating the corresponding changes in aligned coverage, mismatches, and gaps are shown in Fig. S4.

To further validate whether assembly quality improved after BBERT filtering, we used the assembly quality assessment tool MetaQUAST (26) (SI Table S4). At the primary 95% alignment-identity threshold, filtered assemblies retained 93.94% of the bacterial-aligned bases recovered in the unfiltered assemblies, whereas total assembly length decreased by 33.70% and fully unaligned sequence decreased by 94.12%. Consequently, the proportion of assembled sequence aligning to bacterial references increased from 68.27% to 96.73% (SI Table S4). The internal unaligned content between alignment blocks within contigs, which is consistent with structural discrepancies such as large indels or misassembly breaks, fell from 95.4-178.5 to 89.0-155.8 bases per Mbp of bacterial-aligned sequence across the four datasets, corresponding to a pooled reduction of 14.3%. The fraction of bacterial-aligned contigs containing an internal unaligned block decreased from 0.104% to 0.088%, corresponding to a 15.3% relative reduction. Terminal unaligned sequence per bacterial-aligned base decreased to 0.75–0.84 of the unfiltered value across the four datasets, corresponding to a pooled reduction of 20.6%, and, among contigs with a qualifying bacterial alignment, the fraction that was only partially aligned decreased from 16.38% to 15.43%. These trends were consistent across all four datasets and at two alignment-identity thresholds (95% and 80%). Thus, the MetaQUAST analysis corroborates the BLAST findings at the structural level: slightly lower completeness alongside cleaner contig architecture, whereas base-level substitution and small-indel rates, which the BLAST analysis quantified directly, are not captured by the structural decomposition reported by MetaQUAST.

To gauge the potential benefit of BBERT filtering on real-world datasets, we applied BBERT on legume root nodule metagenomes (53). Legume nodules are a useful test case, as the bacterial community is relatively simple and predictable and non-bacterial DNA is rampant due to host contamination. We ran a full metagenomic co-assembly and binning pipeline on raw and BBERT-filtered reads of four nodule metagenomes (see Methods in (53)). The BBERT-filtered co-assembly was 61% smaller (from 577.6 Mbp to 222.4 Mbp), had 57% fewer contigs, and GC content shifted from 40.4% to 58.5%, consistent with the known GC content of legume (∼40%) and *Rhizobiales* (∼62%) genomes, indicating that the vast majority of filtered reads were host-derived. Furthermore, BBERT filtering doubled the assembly’s N50 from 0.9 Kbp to 1.8 Kbp, and halved the assembly run time, while also decreasing peak RAM from 188 GB to 82 GB. Metagenomic bin completeness decreased slightly, consistent with the results from the synthetic data. This caused three low-quality bins from the unfiltered data to drop below the 50% binning threshold in the filtered data, despite their contigs being present. Six bins were recovered by both pipelines. Contamination fell in three bins and rose slightly in the other three. The three contamination decreases occurred in the bins that carried appreciable contamination in the unfiltered assembly (2.7–7.4%, falling to 1.5–3.4%), whereas the three increases occurred in bins that were already essentially pure (0.02–0.76%) and remained so after BBERT filtering (≤1.6%).

The largest single change was for the *Cellvibrio* bin (7.41% to 1.48%). Decomposing both bins by contig GC content shows that the unfiltered bin was bimodal: 5.49 Mbp (58.7%) were low-GC (≤42% GC) similar to the composition of the legume host, and 3.87 Mbp were 48.8% GC, close to that of the *Cellvibrio* sp000263355 species representative. BBERT filtering removed essentially all of the host-derived contigs and left the bacterial contigs intact. Scored against the MIMAG standard (54), BBERT filtering took the assembly from no high-quality draft genomes to one: the BBERT *Bradyrhizobium algeriense* MAG is the only bin in either pipeline carrying a complete 5S, 16S and 23S rRNA set (SI Table S10). All completeness-corrected MAG sizes fall within ∼10% of the expected size for their taxon, with two exceptions (SI Table S10). The unfiltered *Cellvibrio* bin is 2.1-fold the size of its species representative. The *Neorhizobium* bin is undersized in both pipelines; it is an unnamed species, so the genus median is likely an over-estimate of the true target, and this is also the least complete of the six bins (Table 1). In summary, using BBERT before de-novo assembly saves significant costs in compute time and generates truer contigs, while retaining almost the same coverage as unfiltered data.

**Table 1.**
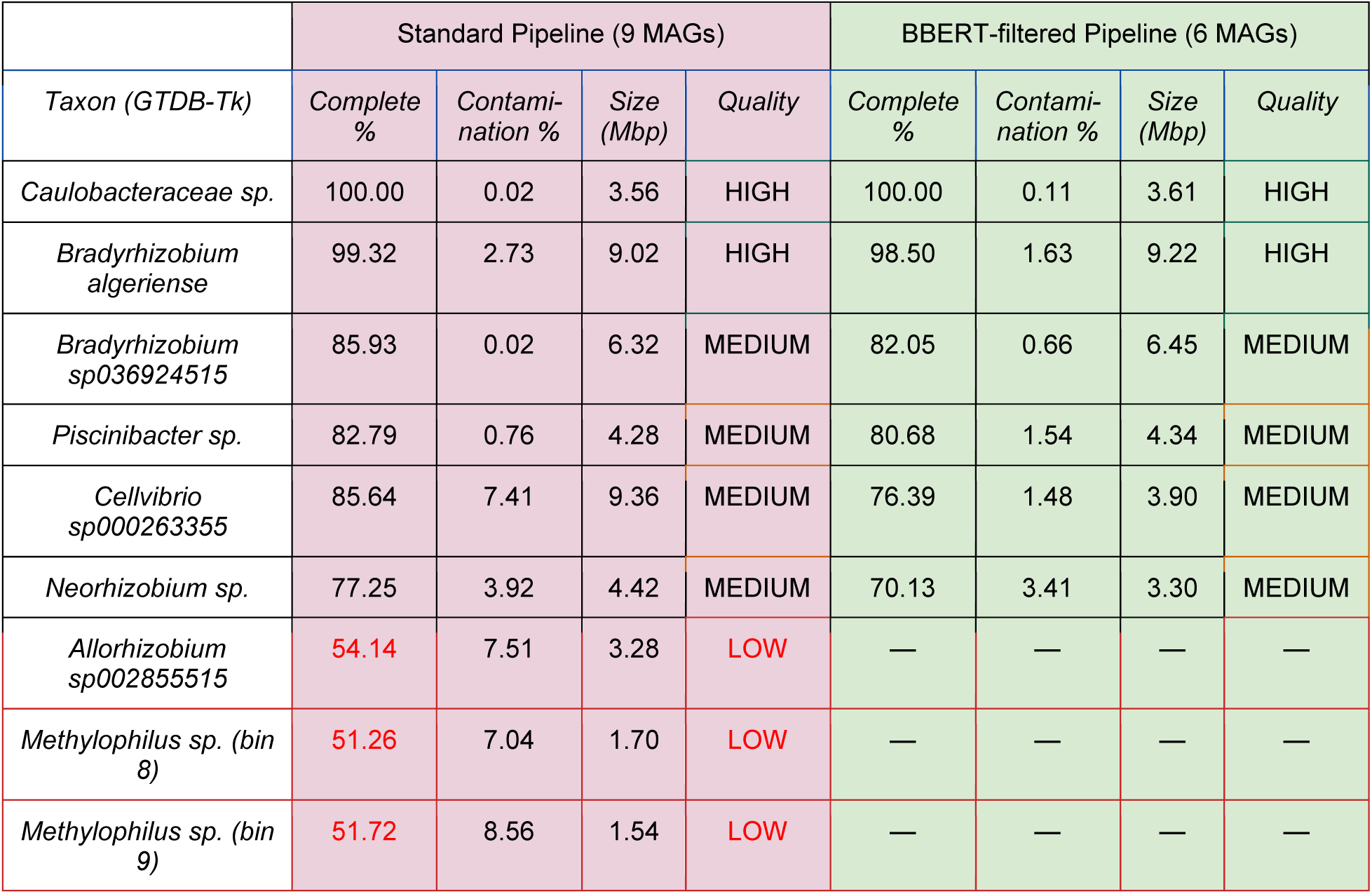
Comparison of quality metrics and taxonomy of MAGs. Completeness and contamination were estimated using CheckM2 (55) (via Binette (56)). GTDB-Tk r220 classification is abbreviated to genus and species where assigned; novel taxa (no species-level assignment) are labelled “sp.”. Quality denotes a completeness/contamination heuristic (HIGH, ≥ 90% complete and < 5% contamination; MEDIUM, ≥ 70%; LOW, ≥ 50%). MIMAG tiers, which additionally requires a complete rRNA operon and ≥ 18 tRNAs, for the same bins are given in SI Table S10.

|  | Standard Pipeline (9 MAGs) |  |  |  | BBERT-filtered Pipeline (6 MAGs) |  |  |  |
| --- | --- | --- | --- | --- | --- | --- | --- | --- |
| *Taxon (GTDB-Tk)* | *Complete %* | *Contamination %* | *Size (Mbp)* | *Quality* | *Complete %* | *Contamination %* | *Size (Mbp)* | *Quality* |
| *Caulobacteraceae sp.* | 100.00 | 0.02 | 3.56 | HIGH | 100.00 | 0.11 | 3.61 | HIGH |
| *Bradyrhizobium algeriense* | 99.32 | 2.73 | 9.02 | HIGH | 98.50 | 1.63 | 9.22 | HIGH |
| *Bradyrhizobium sp036924515* | 85.93 | 0.02 | 6.32 | MEDIUM | 82.05 | 0.66 | 6.45 | MEDIUM |
| *Piscinibacter sp.* | 82.79 | 0.76 | 4.28 | MEDIUM | 80.68 | 1.54 | 4.34 | MEDIUM |
| *Cellvibrio sp000263355* | 85.64 | 7.41 | 9.36 | MEDIUM | 76.39 | 1.48 | 3.90 | MEDIUM |
| *Neorhizobium sp.* | 77.25 | 3.92 | 4.42 | MEDIUM | 70.13 | 3.41 | 3.30 | MEDIUM |
| *Allorhizobium sp002855515* | 54.14 | 7.51 | 3.28 | LOW | — | — | — | — |
| *Methylophilus sp. (bin 8)* | 51.26 | 7.04 | 1.70 | LOW | — | — | — | — |
| *Methylophilus sp. (bin 9)* | 51.72 | 8.56 | 1.54 | LOW | — | — | — | — |

### Analysis of BBERT’s limitations

Finally, to understand how to improve BBERT, we analyzed biases that occur in the bacterial reads that get mis-classified as nonbacterial, i.e. ‘False Negative’ (FN). We tested this by inspecting the results of BBERT on the same four datasets of 512 bacterial genomes used to test the assembly (Fig. 6). We found that BBERT struggles mostly with bacterial non-coding sequences, sometimes mistaking them as nonbacterial (<u>Fig. S5</u>). To identify functional biases in BBERT, we compared Gene Ontology (GO) enrichment between correctly and incorrectly classified reads in both bacterial and eukaryotic test sets. Among eukaryotic reads, BBERT true negatives were enriched for nuclear and multicellular eukaryotic functions, including transcriptional regulation, RNA polymerase II-associated transcription factors, receptor signaling, development, cell differentiation, extracellular/receptor complexes, and cytoskeletal organization. In contrast, eukaryotic false positives were enriched for organellar and bacterial-like bioenergetic functions, including photosynthesis, photosystems, thylakoid/photosynthetic membranes, electron and transmembrane transport, ATP-dependent activity, mitochondrion-associated terms, and small-molecule metabolism. Thus, eukaryotic reads misclassified as bacterial are enriched for organellar and energy-conservation systems with bacterial evolutionary or sequence similarity. The reciprocal bacterial analysis showed that BBERT true positives were enriched for canonical bacterial physiology, including membrane transport, oxidoreductase activity, small-molecule and carbohydrate metabolism, catabolism, ion and organic-anion transport, respiration, and phosphorus metabolism. Bacterial false negatives were enriched for conserved or atypical sequence classes, including DNA binding, DNA polymerase activity, DNA topoisomerase type II regulator and inhibitor activity, integrase and DNA strand-exchange activity, RNA processing including splicing-like annotations, nuclease activity, transposition, toxin-antitoxin, photosystems, organellar ribosomal subunits and plasmid maintenance and partitioning. Together, these results indicate that BBERT efficiently retains ordinary bacterial metabolic and membrane-associated genes, while its main bacterial losses occur in mobile elements, plasmid-associated genes, photosynthetic/organellar-like genes, and conserved informational machinery (<u>Fig. S6</u> and Table S9). Conversely, its main eukaryotic false positives arise from organellar and bioenergetic genes.

Since some of the strongest bacterial false negatives were related to plasmids, we reasoned that BBERT struggles with classifying reads from plasmids, as they are subjected to different evolutionary pressures and dynamics than the bacterial chromosome. It is possible that such plasmid sequences exhibit a distinct DNA syntax that BBERT flags as out-of-distribution. We tested this hypothesis by inspecting the results of BBERT on our mock metagenomes, partitioning reads as originating from one of three groups: chromosomal reads from a genome without plasmids, chromosomal reads from a genome with plasmids, or plasmid reads. To avoid biases resulting from a possible difference in coding vs. noncoding fractions between chromosomes and plasmids, we only considered coding regions. We found a significant difference between false negative rates for reads originating from plasmids vs. reads originating from chromosomes (Fisher test p<1e-300, Odds Ratio >2.2, <u>Fig. S7</u>), with an inconsequential difference between the chromosomal groups (Fisher test p<2.7e-9, Odds Ratio 1.02). This indicates that BBERT flags protein-coding plasmid reads as out-of-distribution more frequently than chromosomal reads, regardless of whether these chromosomal reads come from an organism that has plasmids. Therefore, mobile genetic elements should be considered as a distinct genomic entity with protein-coding DNA following patterns incompatible with chromosomal reads.

To further test whether BBERT misclassifies organellar or conserved eukaryotic sequences as bacterial, we simulated 4.9 million reads from five protein-coding sequence sets: bacteria, mitochondria, chloroplasts and plastids, BUSCO (30) eukaryotic single-copy genes, and random eukaryotic nuclear CDS. BBERT classified bacterial reads most frequently, as expected, but also classified many chloroplast reads as bacterial: 74%, compared with 45% for mitochondria, 31% for eukaryotic single-copy genes, and 28% for random eukaryote nuclear CDS (<u>Fig. S8</u>A). Thus, the main source of bacterial false positives comes from chloroplasts, consistent with their endosymbiotic origin. Gene-level analysis showed that bacterial classification is highest for conserved organellar bioenergetic genes and lower for organellar ribosomal and accessory genes (<u>Fig. S8B-C</u>). Because chloroplasts also encode multiple homologs of the Parks (57, 58) bacterial marker set, chloroplast-rich samples may inflate the BBERT-classified bacterial marker pool. The few CDS that are called bacterial at high rates in eukaryotic nuclear reads are overwhelmingly uncharacterized, lineage-specific hypothetical proteins from reduced-genome parasites (8 of the top 22 from *Enterocytozoon bieneusi*, the others from *Cryptosporidium andersoni*, *Encephalitozoon cuniculi*, *Leishmania braziliensis* and *Chlorella variabilis*). We note that Fig. S8 shows a per-read rate from 52 taxonomically dispersed eukaryotes chosen to span the compositional range, including microsporidia, trypanosomatids and green algae. For genomes of ordinary composition, where most eukaryotic contamination in soil and host-associated metagenomes originates, the bacterial false detection rate is 16%.

## Discussion

Our analysis of a global dataset of 1969 soil metagenomes, using a newly published tool, SingleM (38), estimates that on average, about 32% of soil metagenomic sequence reads are not from prokaryotes (Fig. 1B). The presence of such large quantities of non-bacterial DNA has significant implications for the computation of functional profiles from metagenomic datasets as well as for comparative analyses (39). Yet SingleM, however powerful, is limited to bulk normalization, since it only estimates prokaryotic fractions based on a small set of marker genes, and is unable to classify individual reads. We address the need for high throughput domain-level classification with BBERT, our large-language-model trained to recognize bacterial reads. Remarkably, BBERT yields an estimate of the prokaryotic fraction that closely resembles SingleM’s, while classifying every read as bacterial or non-bacterial (Fig. 4). We tried to assess whether certain branches of the phylogenetic tree are more prone to error and could not identify a statistically significant signal.

BBERT innovates by using LLM technology to assign taxonomy to short, unassembled reads. While theoretically, longer contigs would be much easier to classify (15, 16, 42, 43), such an assembly-based approach is severely limited given current data and computational tools. We showed this by assembling 478 soil metagenomes, where 67% of the reads were not assembled at all (<u>Fig. S1</u>). In comparison, existing tools have been trained on long reads which do not represent the many petabytes of existing short read data (15, 42).

Although binary classification of reads requires only a single score, BBERT’s high-dimensional embedding space captures rich, multidimensional features, including properties beyond those it was trained for. Analysis of BBERT’s embedding space in a mock dataset revealed not only clear domain-level separation but also seven discrete clusters of reads within each genome: one comprising non-coding fragments and six corresponding to the six possible coding reading frames (Fig. 2). This observation suggests substantial potential to enhance existing bioinformatic algorithms by leveraging these embedding-derived structural features. Training classifiers on the embedding space resulted in excellent inference not only of the taxonomic domain, but also whether a read is protein coding and its correct reading frame. Thus, BBERT can improve both the speed and accuracy of many downstream tools. Although BBERT was validated using soil data in this study, it is likely applicable to other metagenomic datasets.

One open question in the metagenomics community is the origin of the so-called “dark matter”, sequence reads with no assigned taxonomy or function. We found that most of the reads that BBERT labeled as nonbacterial could not be annotated by KEGG, whereas the bacterial fraction is mapped successfully by KEGG (Fig. 5A). Conversely, KEGG-unmapped “dark matter” reads are relatively enriched for non-bacterial sequences (Fig. 5B-C). Therefore, it stands to reason that the “dark matter” in soil metagenomes is enriched with eukaryotic genomes, non-coding prokaryote DNA and viral DNA. The presence of large fractions of non-bacterial and unannotated sequences, at varying levels, among datasets, hinders our ability to correctly analyze and compare different habitats to one another. Comparison requires that the data is correctly normalized to sequencing depth. With a large and unknown amount of non-prokaryotic DNA in a sample, it is not possible to know the true sequencing depth of the bacterial genomes in a sample.

Assembly and binning of metagenomic data into putative genomes remains a major challenge in metagenomics. As we have shown here, most metagenomic reads do not assemble at all (<u>Fig. S1</u>). But also, the contigs that are generated de-novo cannot be validated, in the absence of genomic references. Applying BBERT to the data prior to assembly reduces the number of erroneous gaps, and reduces mismatches. This improvement in assembly quality comes with a cost of a 4% loss in alignment coverage but with an almost three-fold speedup in run time, which translates into saved costs (Fig. 6). When we binned the root nodule metagenomic assembly, BBERT-filtered data reduced bin contamination wherever contamination was relatively high, and resolved one bin that the unfiltered pipeline had assembled as a bacterium-host chimera. (Table 1). This opens the possibility of using BBERT to remove host reads of non-sequenced organisms, thus analyzing the bacterial communities of lesser studied eukaryote hosts. An interesting direction to explore is how BBERT could be used to improve de-novo assemblies from metagenomes by adopting a combined approach: assemble the data before and after BBERT filtering. For regions that overlap between the two assemblies, use the assembly of the filtered data, and fill in gaps from the assembly of unfiltered data. BBERT is ready to be used to improve the reliability of metagenomic assemblies.

When sequencing metagenomes, especially from environmental samples, one is bound to encounter a large fraction of genomes not represented in existing references. LLMs are excellent at generalizing classification from partial training data and are therefore well-suited for analyzing environmental metagenomes. The research community has recognized this potential and we have seen an explosion of recent and high quality tools which utilize enormous computational power to derive deep insights into DNA (15–17, 40, 42, 43). A recent and noteworthy related tool is REMME (17), which, like BBERT, was trained on short metagenomic reads. After fine-tuning, REMME achieved good bacterial/non-bacterial classification performance, although it remained below BBERT in our benchmarks. In contrast, REMME’s performance on coding/non-coding classification and reading-frame prediction was substantially lower than BBERT’s (Fig. 3), suggesting that BBERT’s architecture is more suitable for these fine-grained sequence annotation tasks. Here, we showed that by selectively using an LLM on a sharp question, trained with an OOD approach, we can derive deep biological insight from reads that are 100 bases long, requiring only modest computational power, and making our tool accessible to everyone. Thus, LLM technology has the power to transform our understanding of microbial ecology and boost ecological genomics at large.

BBERT’s OOD approach is challenged by the fact that the tree of life includes three domains. This leaves open the question whether to follow the prokaryotic-eukaryotic division and train BBERT on archaea and bacteria together, or to follow the topology of the tree, and use the first split to determine our training strategy. We chose to use the latter and train BBERT only on bacterial genomes. In practice, since archaea and bacteria share many similarities, BBERT appears to classify archaeal reads as within the bacterial distribution. Since archaeal abundance in topsoil is considerably lower than the other two domains (SingleM predicts an average of 2.2±0.06% archaea across the samples we curated), this does not influence the outcome, but caution should be used in habitats where archaea are abundant.

Finally, an examination of BBERT’s limitations also has the potential to reveal hidden biological insight. We found that BBERT struggles with classifying bacterial non-coding DNA and to a lesser extent, mobile genetic elements. Conversely, BBERT tends to classify organelle DNA as bacterial, consistent with its origin. The protein coding language imposes a rigid structure on DNA syntax, and it makes sense that an LLM will be sensitive to this. As we demonstrated above, this sensitivity is quite useful. The fact that BBERT is confused by mobile genetic elements hints at the evolutionary orthogonality of these elements to the host. Lateral gene transfer (LGT) is a major driver of bacterial diversity and evolution, and its history is extremely difficult to retrace (59). The sensitivity of LLM tools to the unique syntax of mobile genetic elements hints at a potential ability of such tools to retrace LGT events with unprecedented sensitivity.

## Acknowledgements

We thank Arnout Schepers and Tenaces Biosciences Ltd. for their support of TS’s independent scientific research outside of his primary professional responsibilities.

## Author contributions

All authors analyzed results and wrote the manuscript. AE, OMF, and TS conceived the idea. DA with TS wrote the BBERT code. MA obtained and curated the metagenomic dataset, as well as the training genome set, and performed the KEGG analyses. GC assisted in bioinformatics analyses. DA and SPM analyzed the embeddings. SPM conducted and analyzed the assembly benchmarks. AE and OMF supervised the research.

## Supplementary data

Supplementary figures and tables S1-S10 are available at NAR online.

## Data availability

BBERT is available to download from www.github.com/AmirErez/BBERT

The code used to evaluate the model and produce the manuscript figures is available from https://github.com/AmirErez/Manuscript_BBERT

The output from the BBERT runs on the 1969 soil metagenomes takes up approximately 3 TB when compressed. It is hosted on our server and is available upon reasonable request.

## Supplementary Figures

**Fig. S1.**
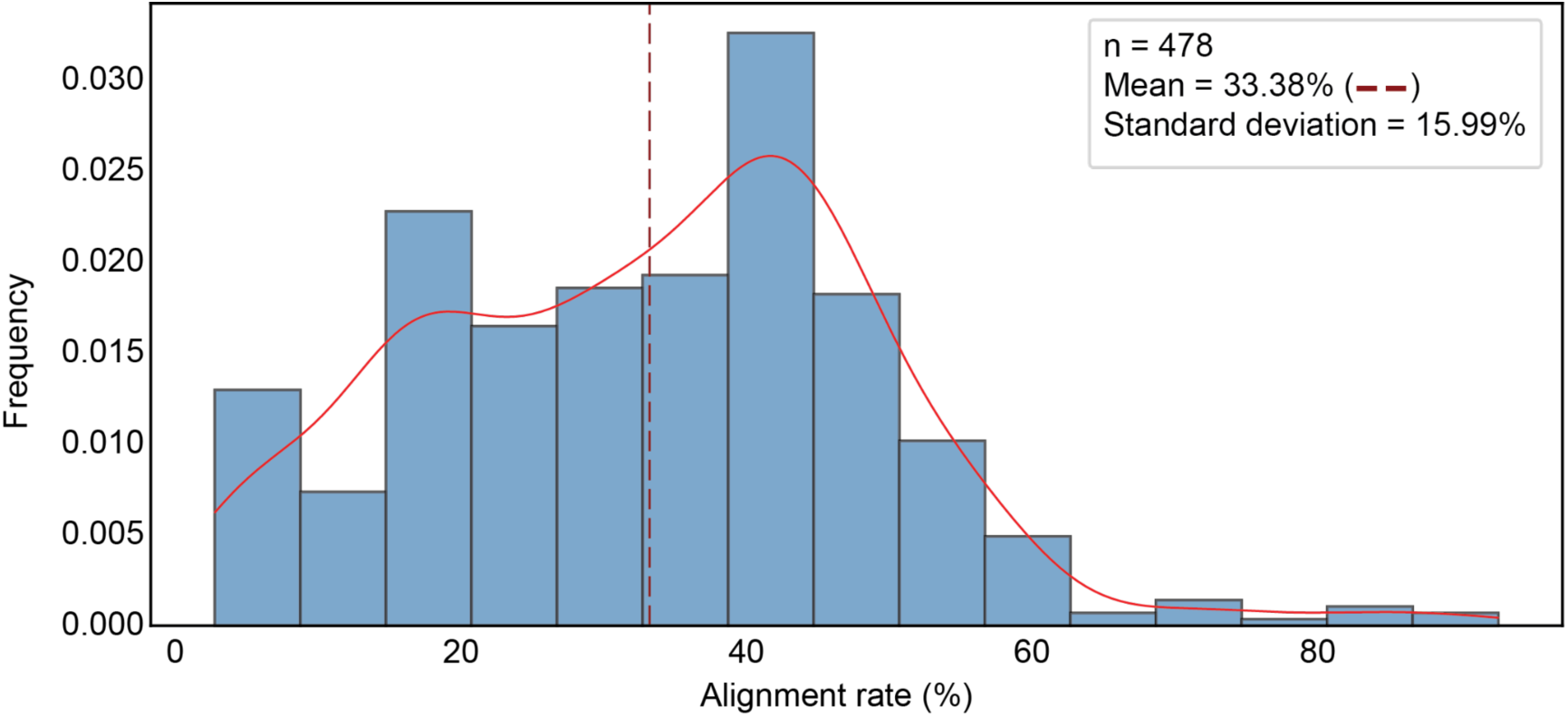
Percent of reads mapped to assemblies of 478 soil metagenomes. The mean alignment rate is 33.38%, meaning that 66.62% of the reads do not align to any contig.

**Fig. S2.**
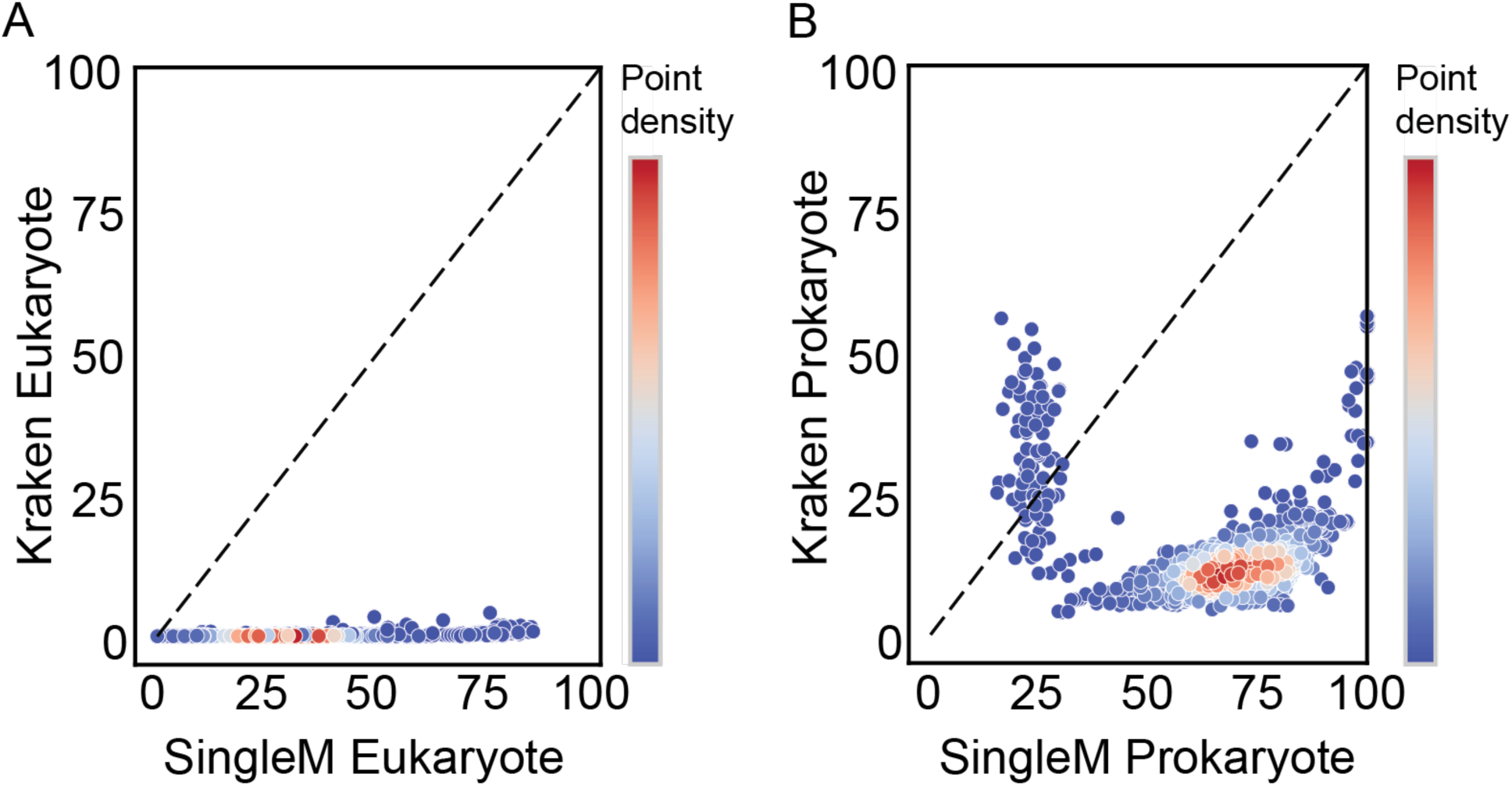
Further evidence for conflicting eukaryote and prokaryote fractions between SingleM and Kraken2’s default database. The imputed domain percent of each sample calculated using SingleM on the *x*-axis and Kraken2 using its default database plus fungi, protozoa, and yeast (PlusPFP, September 2024). **(A)** Imputed eukaryote fraction, where only positively identified eukaryotes are considered for Kraken2. **(B)** The imputed prokaryote percent of each sample.

**Fig. S3.**
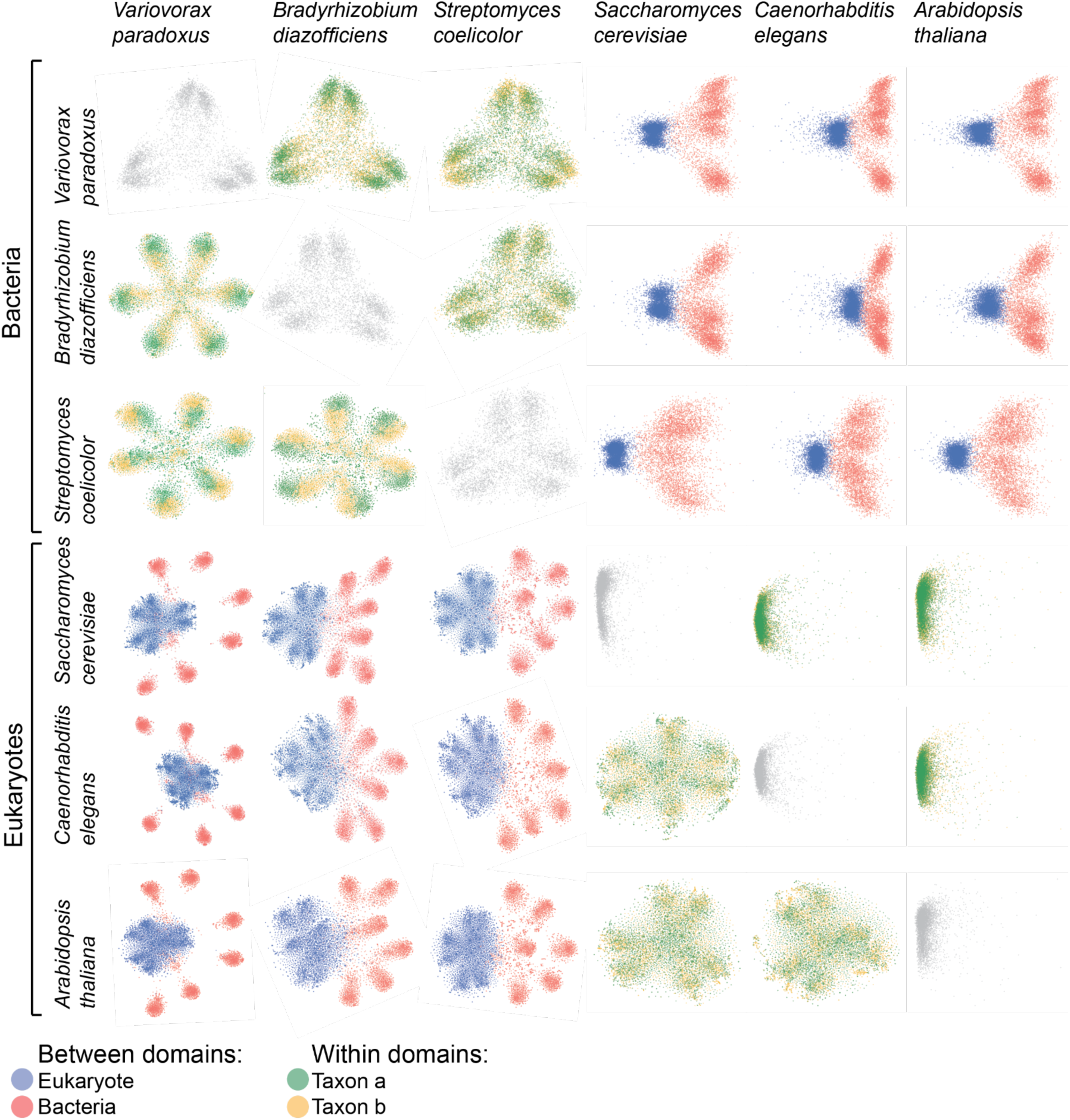
Generalization of BBERT embedding structure across taxa within and between domains. Grid of pairwise taxon comparisons derived from per-read BBERT embeddings. Off-diagonal panels show joint embeddings of the two indicated taxa: the lower triangle uses t-SNE, and the upper triangle shows PCA (PC1 vs. PC2). Diagonal panels display single-taxon PCA to highlight within-taxon substructure. Across pairs, both methods robustly separate taxa, including across domains, while single-taxon views reveal finer organization consistent with content-based features (Fig. 2).

**Fig. S4.**
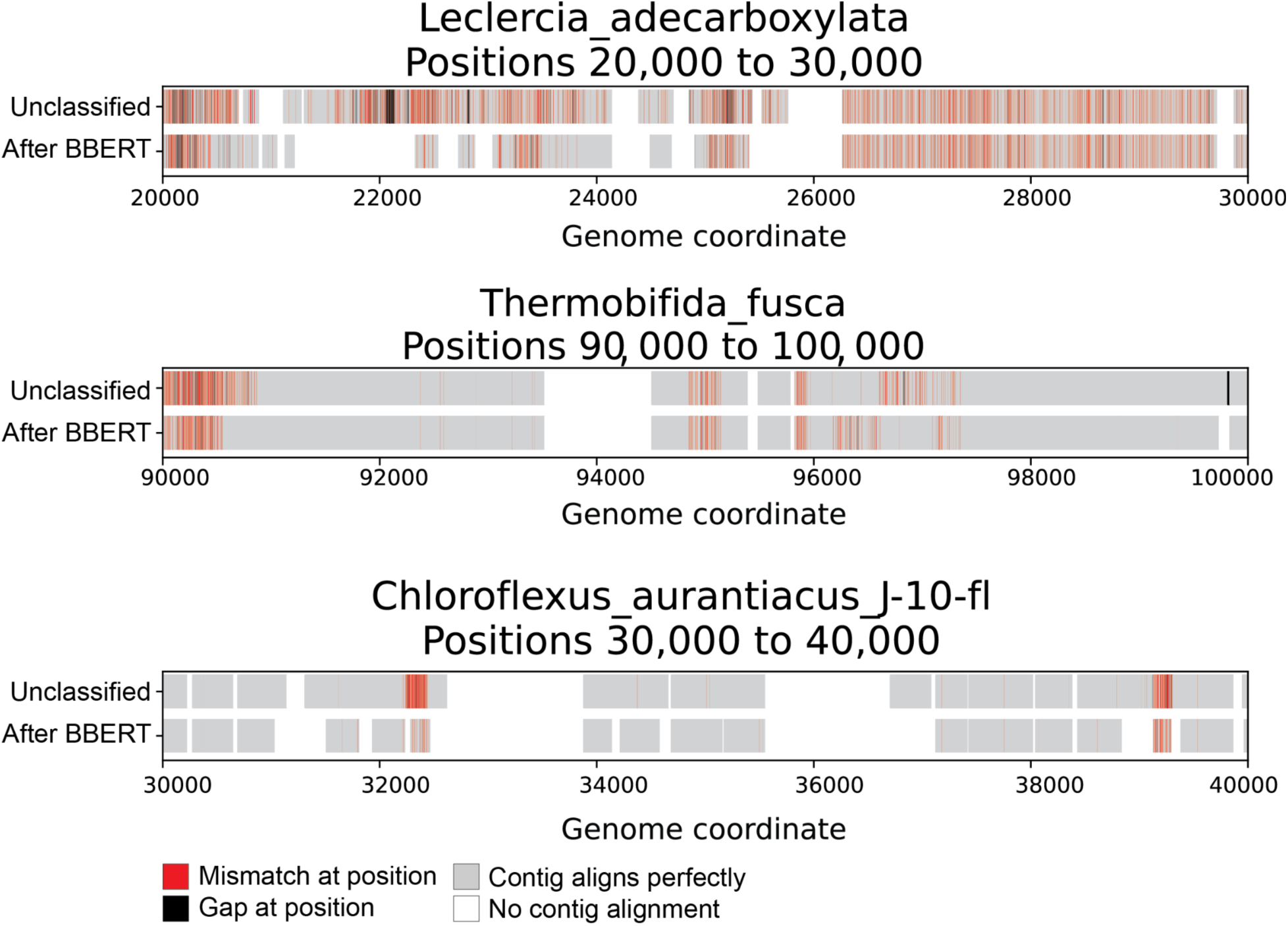
Comparison of de-novo assembly errors before and after BBERT filtering. Reads were assembled with metaSPAdes^25^ using either unfiltered input or BBERT-filtered (bacterial) reads. Resulting contigs were aligned to the known bacterial source genomes used to generate the mixtures (see Methods), enabling per-base annotation of mismatches and gaps and identification of mis-assemblies. Shown are representative loci illustrating some outcomes: unfiltered data produces chimeric joins or dense local errors, whereas BBERT-filtered data assembles the region correctly or refrains from assembling (avoiding false contigs); in a minority of cases, the unfiltered assembly is correct while filtering prevents assembly, consistent with reduced coverage after removal of non-bacterial reads; some regions remain challenging under both conditions. Aggregate improvements in assembly quality and runtime are quantified in Fig. 6.

**Figure S5.**
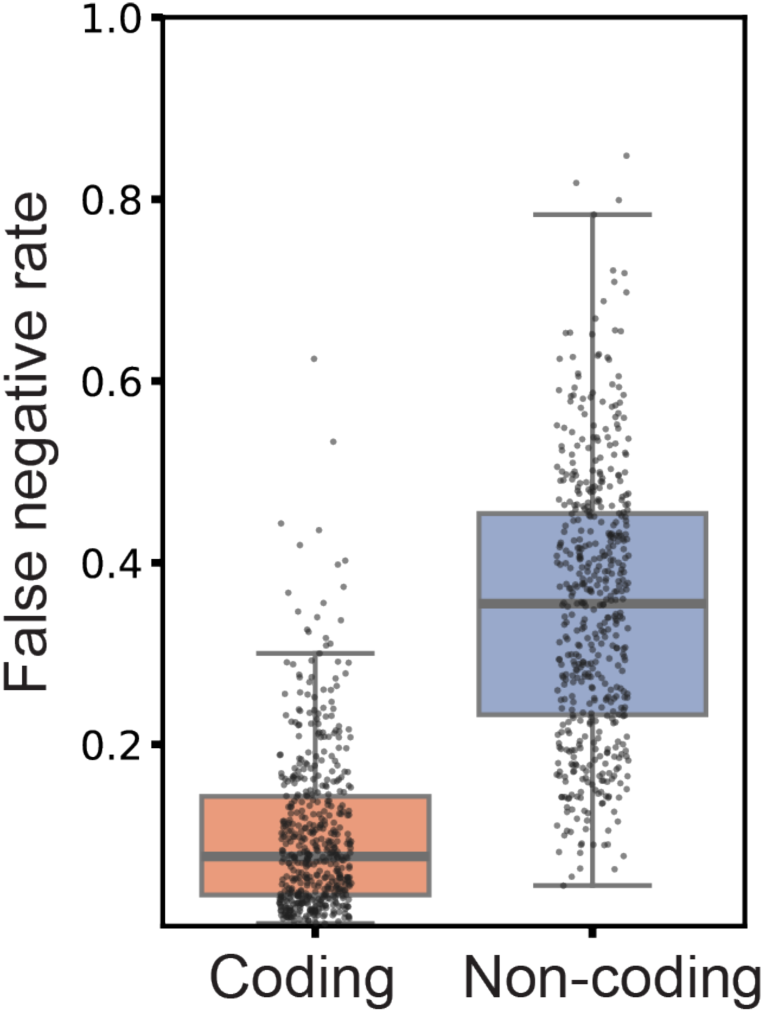
Comparing BBERT results on coding vs. noncoding bacterial reads. We used the four datasets of 128 bacteria genera each described in Methods Section “**Reduced assembly errors and run-time acceleration**” (∼140K reads).

**Figure S6.**
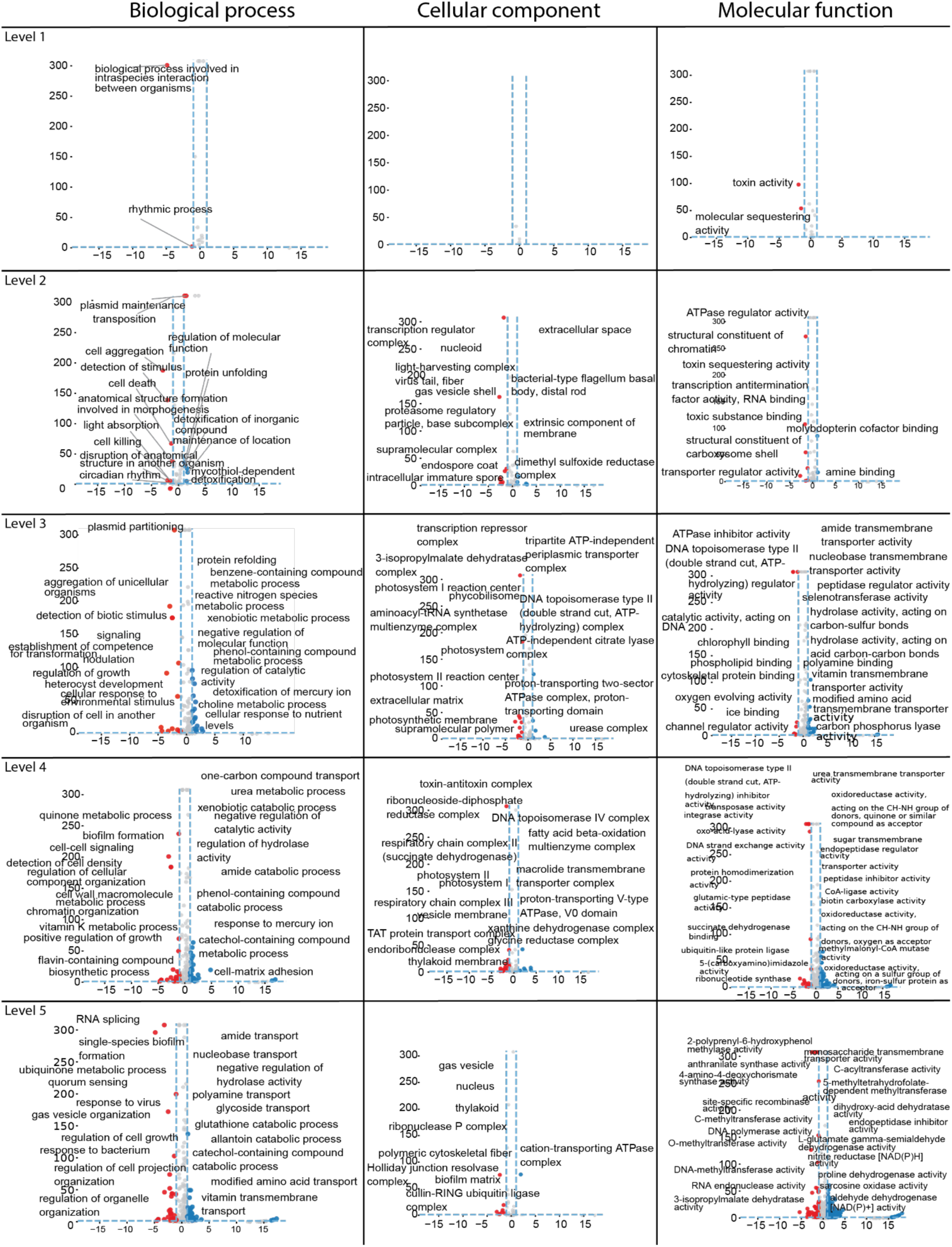
Gene Ontology analysis of terms enriched for False Negative and True Positive. Volcano plots for bacterial classification enrichment in true positive (blue) and false negative (red). The top 10 most significant GO terms are labeled at five GO levels (Table S9). The GO annotation process is described in the Methods section.

**Figure S7.**
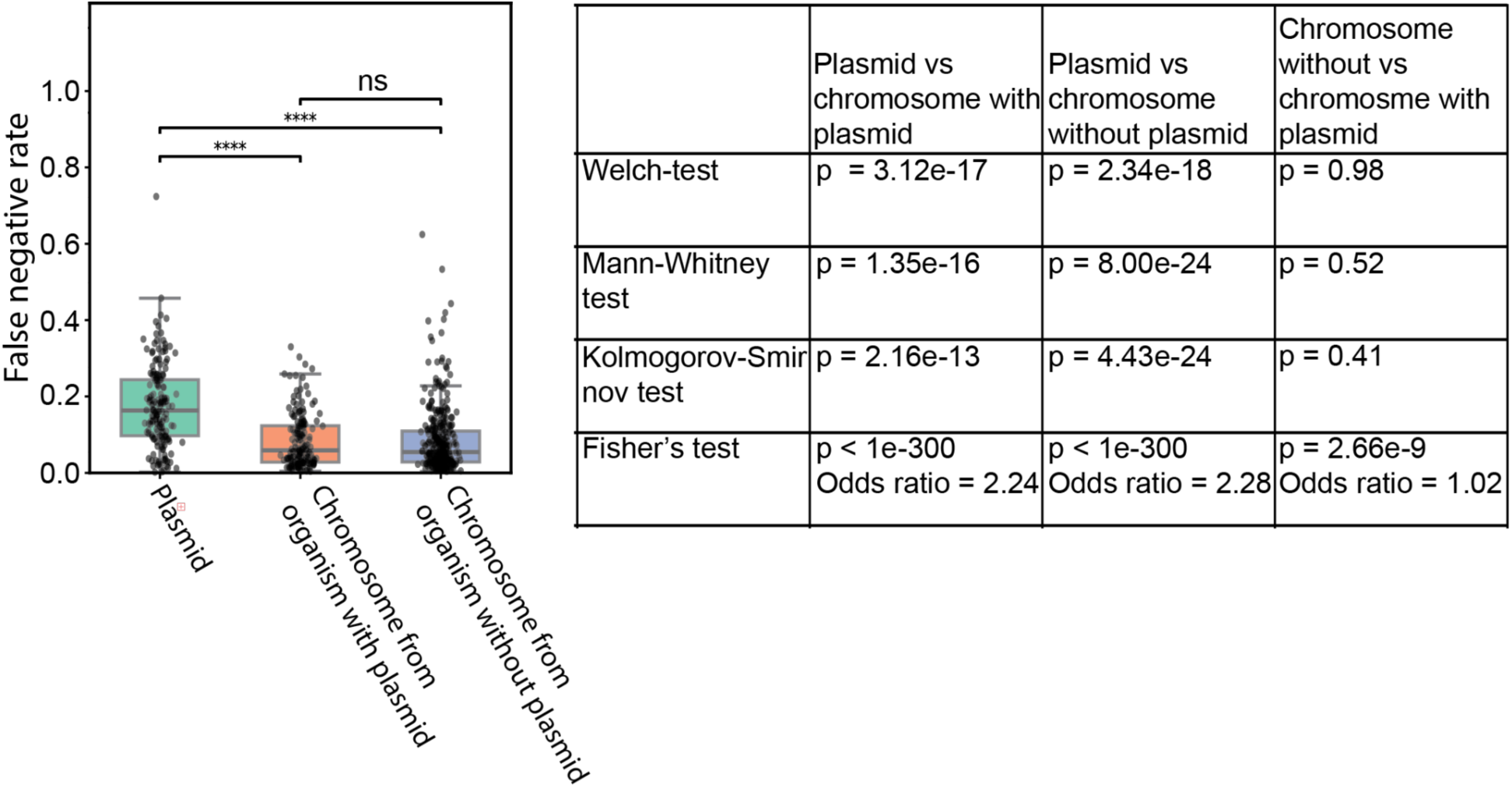
Comparing BBERT results on plasmid vs. non-plasmid bacterial reads. For each read, we determined whether it originated from a chromosome or a plasmid. Organisms were divided into two groups: those with at least one plasmid and those without any plasmids. Plasmids were identified based on the NCBI genome headers. We labeled the groups as follows: Plasmid, Chromosome in organisms with plasmids (ChromP), and Chromosome in organisms without plasmids (ChromN). We calculated the FN rates for each organism; statistical tests were applied to assess differences between the groups. Based on the Shapiro-Wilk normality test (Plasmid=1.574e-13, ChromP=1.755e-04, ChromN=1.522e-03), we performed group-to-group comparisons using the t-test, Mann-Whitney, and Kolmogorov-Smirnov tests. The results show a strong difference between the plasmid and non-plasmid groups, with no significant difference between chromosomal groups (Fisher test shows an Odds Ratio of 1.06). This indicates that BBERT tends to misclassify plasmid reads much more frequently than chromosomal reads, regardless of whether these chromosomal reads come from an organism that has plasmids. The significance shown is Mann-Whitney test results.

**Fig. S8.**
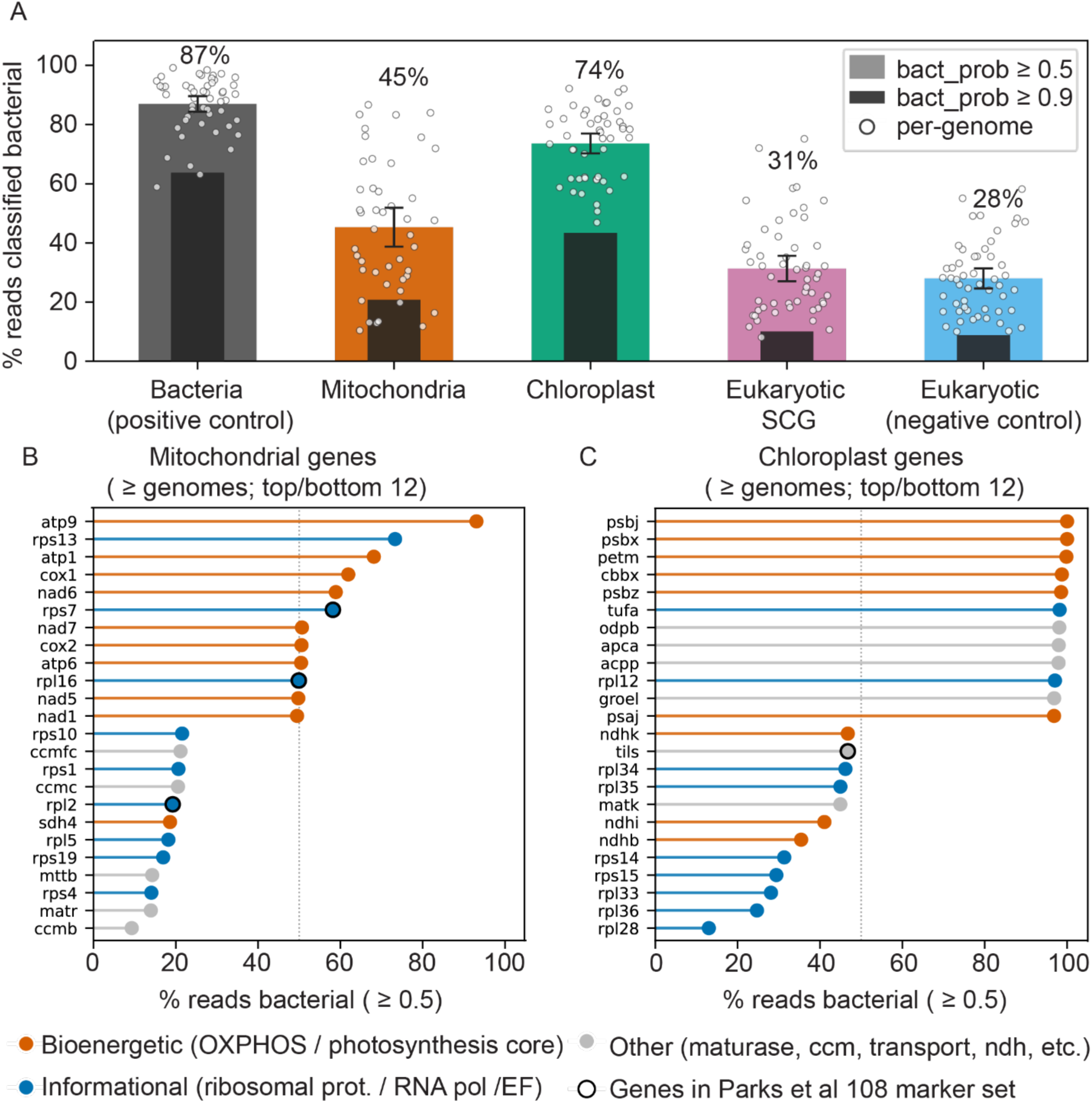
Mitochondrial, chloroplast and eukaryotic single-copy genes classified as bacterial by BBERT. **(A)** Percentage of reads classified bacterial per category (mean across 50 genomes ± 95 % CI; grey bars, bact_prob ≥ 0.5; black overlay, ≥ 0.9; open circles, per-genome values). **(B-C)** Per-gene bacterial-classification rate (bact_prob ≥ 0.5) for **(B)** mitochondrial and **(C)** chloroplast genes present in at least 3 genomes (the 12 most- and 12 least-bacterial shown). Genes are colored by functional class: orange: bioenergetic (photosynthesis core); blue: informational (ribosomal proteins, RNA polymerase, translation factors); grey: other (maturases, cytochrome-c maturation, transport, NDH-like, ORFs). Ringed markers belong to the 108-gene Parks single-copy marker gene set used by SingleM. The conserved, endosymbiont-derived bioenergetic core classifies as bacterial most often, whereas the divergent organellar ribosomal proteins and organelle-specific accessory machinery classify as bacterial least often.

